# Automated live-cell single-molecule tracking in enteroid monolayers reveals transcription factor dynamics probing lineage-determining function

**DOI:** 10.1101/2024.04.04.587889

**Authors:** Nike Walther, Sathvik Anantakrishnan, Gina M. Dailey, Robert Tjian, Xavier Darzacq

## Abstract

Lineage transcription factors (TFs) provide one regulatory level of differentiation crucial for the generation and maintenance of healthy tissues. To probe TF function by measuring their dynamics during adult intestinal homeostasis, we established HILO-illumination-based live-cell single-molecule tracking (SMT) in mouse small intestinal enteroid monolayers recapitulating tissue differentiation hierarchies *in vitro*. To increase the throughput, capture cellular features, and correlate morphological characteristics with diffusion parameters, we developed an automated imaging and analysis pipeline, broadly applicable to 2D culture systems. Studying two absorptive lineage-determining TFs, we find an expression level-independent contrasting diffusive behavior: While Hes1, key determinant of absorptive lineage commitment, displays a large cell-to-cell variability and an average fraction of DNA-bound molecules of ∼32%, Hnf4g, conferring enterocyte identity, exhibits more uniform dynamics and a bound fraction of ∼56%. Our results suggest that TF diffusive behavior can indicate the progression of differentiation and modulate early *versus* late differentiation within a lineage.

**Highlights:**

- Automated live-cell single-molecule tracking records hundreds of cells in enteroid monolayers
- Cellular diffusion clustering and morphological feature correlation reveals subpopulations
- Transcription factor dynamics regulate differentiation independent of expression level
- Hes1 and Hnf4g display contrasting dynamics assisting early *vs.* late absorptive differentiation

## Introduction

During adult tissue homeostasis, stem cells continuously give rise to the various cell types of a tissue. Stem cell differentiation is characterized by a rewiring of the cellular gene expression program. Downstream of signaling pathways, lineage-determining transcription factors (TFs) regulate lineage commitment and further differentiation to the mature cell types comprising a tissue^1^. TFs induce or repress the expression of their target genes by recruiting co-regulators and binding to *cis*-regulatory elements^2^. Key TFs conferring lineage commitment or cell fate decisions have been identified in various tissues and their misexpression is associated with disease^3,4^. However, it remains poorly understood whether the molecular dynamics of cell fate-determining TFs are correlated with differentiation and how they regulate early lineage commitment and later cell fate decisions along the same differentiation pathway of stem cells to mature cell types. Filling this knowledge gap requires the combination of live cell-compatible methods allowing to probe TF dynamics at the single-molecule level with model systems recapitulating differentiation trajectories found *in vivo*.

Such a system has been derived from the mammalian intestine, which is the fastest renewing organ and consists of intestinal stem cell (ISC)-containing crypts and villi composed of mature cell types. A spatial differentiation hierarchy along the crypt-villus axis guides directional movement of differentiating cells: ISCs divide to either self-renew or give rise to lineage-committed proliferative absorptive or postmitotic secretory progenitors in the transit-amplifying zone (TAZ), which, upon further maturation and final cell fate commitment, move towards villi^5,6^ (Fig. 1A). These central features of the gut are recapitulated in *in vitro* tissue models: Mouse small intestinal organoids (mSIOs; enteroids) are 3D budding structures with ISC-containing crypts at their tips and differentiating ISCs moving inward^7,8^ (Fig. 1A). Such organoid models are amenable to live-cell imaging and have recently allowed for revolutionary studies of cellular behavior within multicellular tissue-resembling systems^9–13^. In a dimensional reduction and simplification, mSIOs can also be grown in 2D as enteroid monolayer cultures (EMCs)^14–16^, which equally consist of all intestinal epithelial cell types and display the spatial differentiation hierarchy observed *in vivo*: ISCs in proliferative centers move outward along a radial differentiation trajectory^14^ (Fig. 1A).

**Figure 1:**
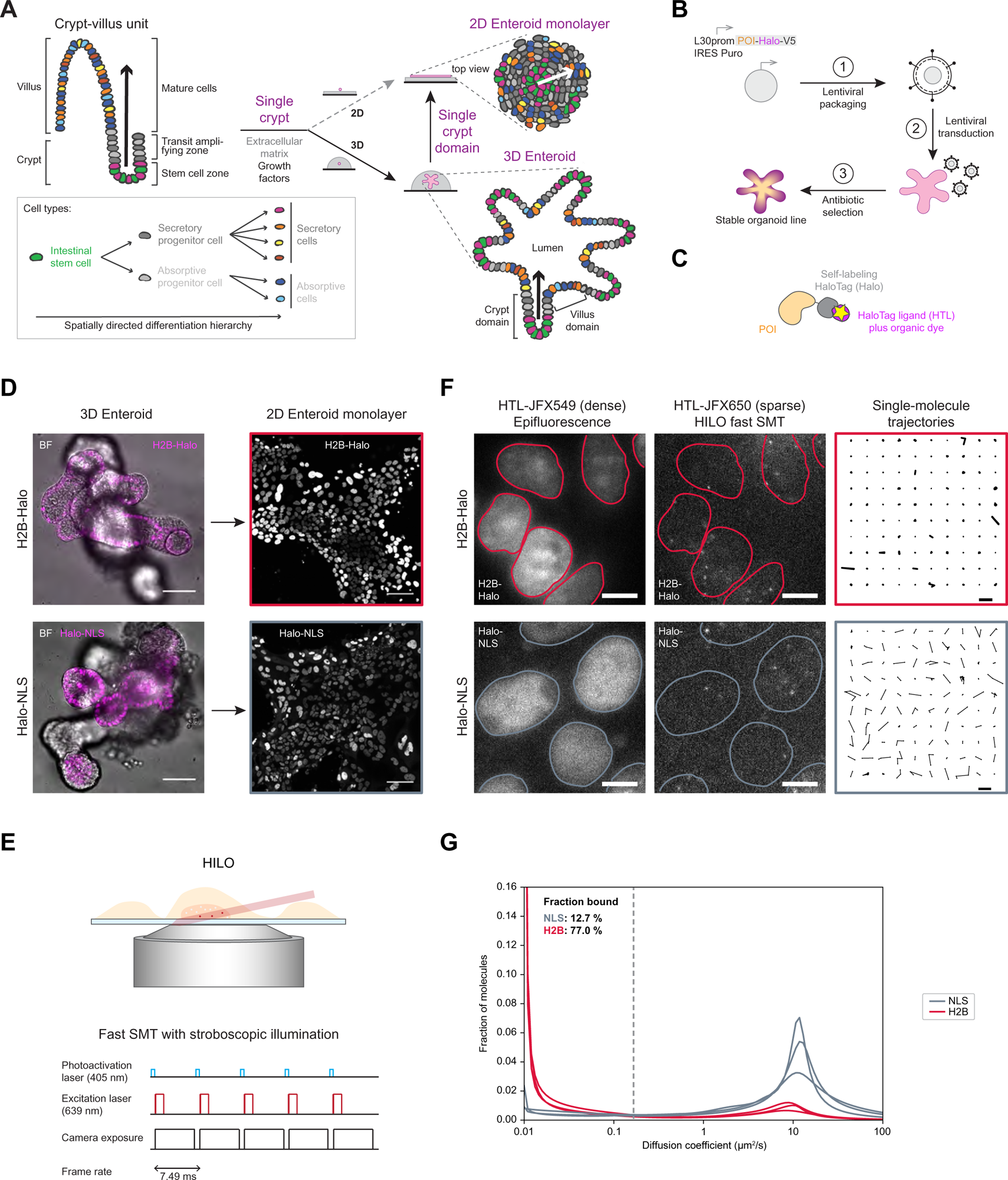
Fast live-cell single-molecule tracking (SMT) in 2D enteroid monolayer cultures (EMCs) determines the diffusive behavior of Halo-tagged proteins in an intestinal differentiation model. **(A)** The differentiation of intestinal stem cells (ISCs; green) to mature absorptive (blue) and secretory (yellow-red) cell types underlies a spatial differentiation hierarchy (black arrow) from ISC-containing crypts over the transit amplifying zone (TAZ) with absorptive/secretory (light/dark gray) progenitor cells towards villi composed of differentiated cell types. Mouse small intestinal crypts embedded into an extracellular matrix give rise to 3D enteroids (bottom right). Cultured in 2D, these organoids form EMCs (top right). Enteroids display the same spatial differentiation hierarchy in 2D from proliferative centers outward (top; white arrow) as in 3D from ISC-containing tips of buds inward (bottom; black arrow). **(B)** Generation of organoid lines stably overexpressing HaloTag-V5 (magenta-black) fusions of the protein of interest (POI; orange) through 1) lentiviral packaging and lentivirus production, 2) transduction of 3D enteroids (rose), and 3) antibiotic selection of transgene expressing enteroids (orange-magenta). **(C)** Covalent HaloTag (gray) labeling with HaloTag ligands (HTLs) coupled to bright and photostable dyes (magenta) confers fluorescence (yellow asterisk), enabling POI detection in live-cell imaging applications. **(D)** Stable enteroid lines expressing H2B-Halo (top) or Halo-NLS (nuclear localization sequence; bottom). Confocal imaging of 3D mSIOs (left; POI (magenta); brightfield (BF; gray)) and corresponding 2D EMCs (right; POI (gray) 5 d post-seeding. Scale bars: 50 μm. **(E)** Principle of HILO illumination (top) for fast SMT of POI-Halo using stroboscopic illumination (bottom): A laser beam (red) is sent obliquely through sparsely labeled cells (orange) close to the glass bottom (blue) of a culture dish (top). Single-molecule movies consist of thousands of imaging frames recorded at a frame interval of a few milliseconds, whereby the excitation pulses (red) are shorter than the camera exposure time (black) to reduce motion blur. Photoactivation pulses (blue) to reactivate dark molecules are spaced in between detection windows. **(F)** Double labeling of POI-Halo with two different HTLs allows bulk labeling for nuclear detection and segmentation (left; red/gray masks) followed by HILO-based SMT (middle) resulting in single-molecule trajectories (right). Chromatin-bound H2B (top; red) and freely diffusing NLS (bottom; gray) served as slowly and fast diffusing controls. The same FOV for densely (left) and sparsely (middle) labeled POIs is shown for one time point of a SMT movie. 100 randomly selected single-molecule trajectories from the same SMT movie are shown (right). Scale bars: 5 μm (left, middle); 1 μm (right). **(G)** Mean diffusion spectra for 3 SMT experiments each for H2B (red) and NLS (gray). Bootstrap analysis of combined experiments with *n*=53,35,39 cells for H2B and *n*=16,42,34 cells for NLS determined a mean fraction bound (*D*<0.15μm^2^/s) of 77% (95% confidence interval (CI): 74.5-79.6%) and 12.7% (95% CI: 10.3-16.5%), respectively.

Differentiated absorptive enterocytes are the most abundant cell type of the gut. They are located at villi, where they function in nutrient absorption. Exposed to harsh mechanical and chemical conditions, enterocytes are replenished every three to five days during adult intestinal homeostasis^5,6^. Hes1 is the key TF regulating commitment to the absorptive lineage^17^, while hepatocyte nuclear factor 4 gamma (Hnf4g) was identified as a TF conferring enterocyte identity^18,19^.

Hes1 is expressed in proliferative ISC and progenitor cells and acts as a transcriptional repressor of the *Atoh1* and *Neurog3* genes^17,20–27^, regulating the differentiation of secretory cells^20,21^. The *Hes1* gene is a downstream target of Notch signaling, which dictates an alternating pattern between secretory and absorptive cells through lateral inhibition^28^. Constitutive overexpression of Hes1 in mice results in the absence of the secretory lineage, whereas its gene knock-out increases the fraction of secretory cells^17,28^. Hes1 is a basic helix-loop-helix TF binding to N-box DNA motifs. It undergoes self-regulation through a negative feedback loop whereby Hes1 dimers bind to the Hes1 promoter, repressing *Hes1* gene expression^29^.

Hnf4g^30,31^ has important functions for digestive physiology and intestinal development^18,32,33^. It is expressed specifically in enterocytes^19^ and was identified as a major driver of enterocyte differentiation^18^. Hnf4g^-/-^ mice are viable with minor gastrointestinal defects^34^. In organoids, a shift towards secretory cell types was detected in the absence of Hnf4g. However, a small enterocyte population remained, suggesting a partial redundance of enterocyte fate determination by other TFs^18^. Notably, Hnf4g can form both homo- and heterodimers with its paralog Hnf4a^35^.

While early cell fate commitment in the intestine is plastic to enable rapid reversal and dedifferentiation upon stem cell damage, late cell fate decisions are irreversible to maintain the differentiated cell types performing critical functions of the gut. This early permissiveness was demonstrated on the chromatin level^36,37^. However, it has not been investigated how the molecular dynamics of lineage TFs might change during differentiation to support early *versus* late cell fate decisions during the same differentiation pathway. Consistent with their function during early *versus* late absorptive differentiation to enterocytes, Hes1 RNA expression decreases from ISCs to mature enterocytes while for Hnf4g it increases^38^. Hence, Hes1 and Hnf4g provide a *bona fide* pair of early and late absorptive lineage TFs to comparatively study their diffusion dynamics during differentiation.

Live-cell single-molecule tracking (SMT) allows the direct measurement of the dynamics of individual protein molecules and thus can resolve differently behaving subpopulations within the same cell. In contrast, other fluorescence-based live-cell microscopy methods determine bulk protein dynamics^39^. In SMT, individual proteins are detected, localized, and tracked over thousands of imaging frames at an interval of only a few milliseconds (fast SMT). Diffusion spectra derived from single-molecule trajectories provide the diffusion coefficients of differentially mobile subpopulations and the fraction of immobile molecules, which, in the case of TFs, are largely chromatin-bound (fraction bound)^40,41^.

Over the last decade, SMT emerged as a powerful technique to probe TF dynamics^42–46^. Hence, TF dynamics have been studied extensively by highly inclined and laminated optical sheet (HILO)-based SMT in 2D cell culture systems, such as immortalized cancer cells^42,45,47,48^ or embryonic stem cells^43,46^. However, academic-scale SMT has been limited in throughput and the information gained has typically been restricted to parameters describing TF diffusion. While first studies have compared two distinct undifferentiated and differentiated states^44,49^, they were obtained through directed differentiation protocols and TF dynamics have thus been measured at different time points outside the multicellular context of a differentiating tissue model. In contrast, live-cell SMT has been performed in embryos of various species^50–53^. However, SMT in such thicker specimen requires advanced microscopy technology, such as lattice light-sheet microscopy^54^, which is more difficult to operate and analyze, limited in throughput, difficult to automate, and generates large amounts of data.

To address the current knowledge gap of whether the diffusive behavior of lineage TFs changes during differentiation, we focused on the absorptive lineage by studying Hes1 and Hnf4g. To enable their dynamic imaging, we generated mSIO lines stably expressing HaloTag (Halo) fusion proteins and established live-cell highly inclined and laminated optical sheet (HILO)-based SMT in derived 2D EMCs. To increase the throughput, we developed an automated pipeline for imaging, data processing, and protein diffusion analysis, which enables cellular feature extraction and correlation with diffusive protein behavior, as well as diffusion-based protein cluster analysis at the single-cell level. Imaging hundreds of cells by automated live-cell SMT allowed us for the first time to capture different cell states and types at the same time in a complex differentiating system and to break it down into subpopulations with similar cellular morphological and diffusive TF characteristics indicative of the progression along a differentiation trajectory. Finally, we implemented automated proximity-assisted photoactivation (PAPA)-SMT^55^ experiments to probe for TF self-interaction. We propose a data-driven model of differential diffusive behavior of TFs acting early *versus* late during intestinal absorptive differentiation, supporting early permissive *versus* late irreversible cell fate decisions.

### Design

Combining the two worlds of a tissue-recapitulating *in vitro* model system with the ease of HILO-based SMT, we established SMT in 2D EMCs to study the dynamics of lineage TFs at multiple differentiation states within the same sample. Consisting of various cell types and states along a differentiation trajectory, sampling a magnitude more cells in such monolayers is required to capture this heterogeneity. To 1) increase the throughput and ease of data acquisition, 2) record additional cellular features, and 3) directly plug the acquired SMT data into an analysis workflow, we developed an automated pipeline for HILO-based SMT, data processing, and analysis that is also compatible with PAPA-SMT for detecting molecular interactions. We further implemented a hierarchical clustering approach based on single-cell diffusion spectra for determining the similarities between TFs and the degree of cellular heterogeneity of TF diffusive behavior. Correlating TF diffusion dynamics with features of cellular morphology will help move SMT towards extracting cell states and types, similar to single-cell transcriptomics methods^56^, as we demonstrate for intestinal absorptive differentiation by investigating the lineage TFs Hes1 and Hnf4g.

Our developed SMT approach bridges scales from simplified 2D *in vitro* differentiation-recapitulating monolayer cultures to single molecules and is compatible with high-throughput assays and perturbations. We therefore anticipate a broad application potential of our developed technology to other 2D cell culture systems to study the molecular mechanisms of a variety of biological processes.

## Results

### Fast live-cell SMT in 2D enteroid monolayers determines the diffusive behavior of HaloTag fusion proteins

To study the diffusive behavior of TFs with lineage-determining roles in the gut, we established HILO-based live-cell SMT in 2D EMCs^14,15^ derived from 3D mSIOs, which consist of all cell types and display the spatial differentiation hierarchy found in the small intestinal epithelium^14^ (Fig. 1A). To enable the visualization of proteins of interest (POIs) by live-cell microscopy, we delivered POI-Halo transgenes lentivirally into 3D organoids to generate stable lines expressing these transgenes from a weak ubiquitous L30 promoter resulting in a slight overexpression in addition to the untagged endogenous protein (Fig. 1B). Covalent labeling of the HaloTag with a HaloTag ligand (HTL) coupled to bright and photostable dyes^57^ (Fig. 1C) allowed for the confirmation of transgene expression in 3D organoids and 2D EMCs (Fig. 1D) and was essential for SMT (Fig. 1E,F). Here, we used HILO-based live-cell fast SMT in combination with a stroboscopic illumination scheme (Fig. 1E) to reduce motion blur^40,46^ on a total internal reflection fluorescence (TIRF) microscope. This allowed for the detection, localization and tracking of thousands of single molecules corresponding to sparsely labeled POI-Halo. Co-staining with a different higher-concentrated fluorophore-coupled HTL led to bulk labeling of nucleus-localizing TFs and enabled nuclear segmentation for the removal of single-molecule trajectories outside of the nucleus (Fig. 1F). We benchmarked our SMT approach in 2D EMCs using Halo-tagged chromatin-bound histone H2B and freely diffusing nuclear localization sequence (NLS) and determined the dynamic range for the fraction of bound molecules to be between 12.7% for NLS and 77.0% for H2B (Fig. 1G; Fig. S1).

### Generation of organoids and derivation of 2D enteroid monolayers expressing Hes1 and Hnf4g for probing TF function by SMT-based diffusion analysis

TFs move around in the nucleus, search for their specific DNA target motifs, and, upon binding, regulate the expression of their target genes^2^. These action steps are reflected by various diffusive states (Fig. 2A). Thus, measuring diffusive protein behavior by SMT allows to probe TF function by reading out diffusive populations as peaks in the diffusion spectrum and the fraction of immobile DNA-bound TF molecules^40^ (Fig. 2A).

**Figure 2:**
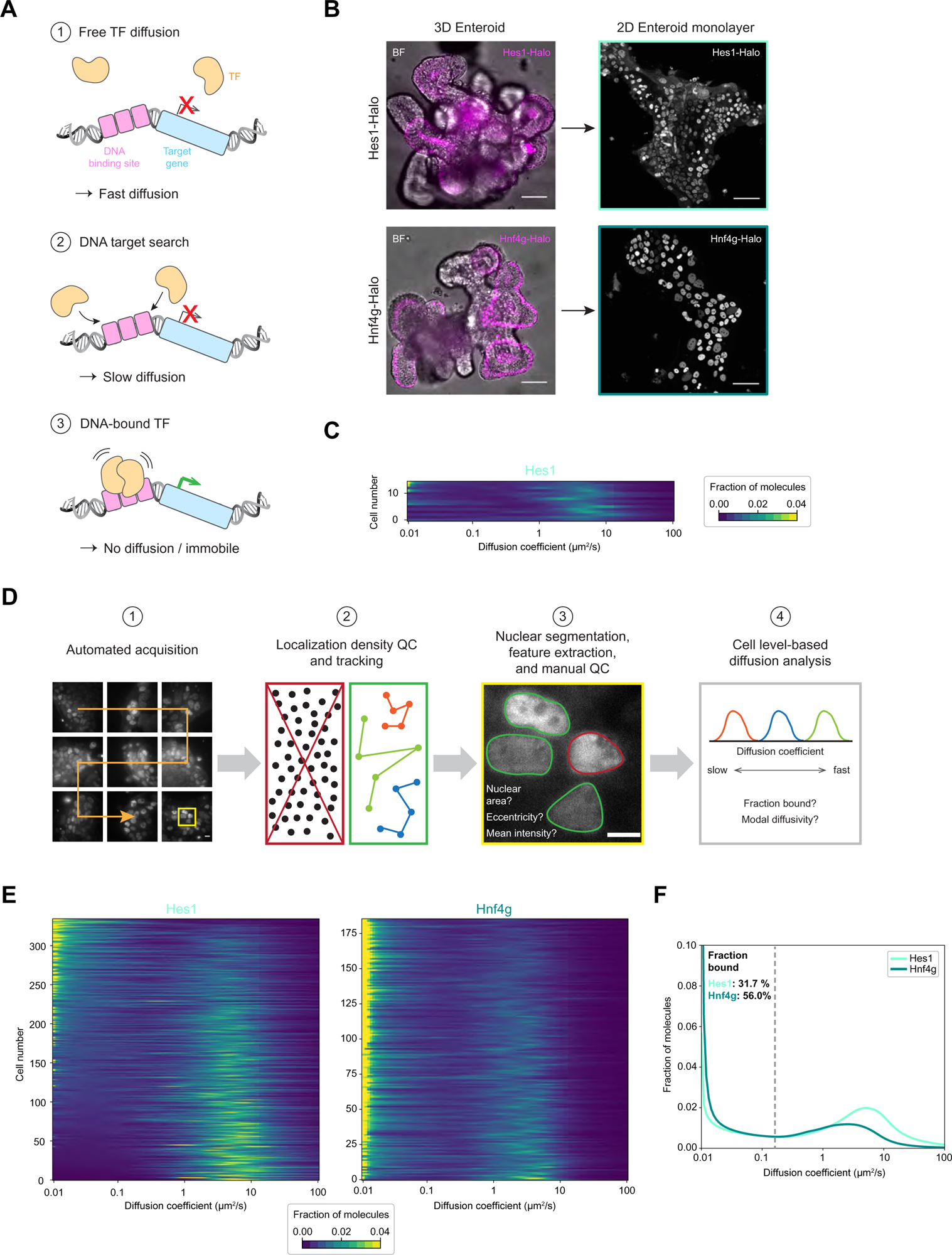
Automated SMT reveals the diffusive behavior of the absorptive transcription factors (TFs) Hes1 and Hnf4g across hundreds of cells in differentiating 2D EMCs. **(A)** Probing nuclear TF diffusion dynamics indicative of TF function: 1) In a non-DNA-bound state, a TF (orange) diffuses freely in the nucleus. 2) Upon encountering DNA (gray), a TF slows down in diffusion while searching for its DNA target site (magenta). 3) TF binding to a DNA target site results in a non-diffusive, immobile state necessary for executing TF function (green arrow) on target gene expression (blue). **(B)** Stable mSIOs expressing Hes1-Halo (top) or Hnf4g-Halo (bottom). Confocal imaging of 3D mSIOs (left; TF (magenta); BF (gray)) and corresponding 2D EMCs (right; TF (gray)) 5 d post-seeding. Scale bars: 50 μm. **(C)** Single-cell diffusion heatmap of a manual fast SMT experiment in Hes1-Halo 2D EMCs. Cells are ordered according to increasing fraction bound from bottom to top (*n*=15 cells). **(D)** Automated imaging and analysis pipeline for HILO-based SMT to measure POI diffusion dynamics in hundreds of cells: 1) Automated imaging in a user-defined grid size (orange arrow). At each position, an overview image is taken in the densely labeled POI channel (here: Hes1-Halo) in epifluorescence mode. Facilitated through the nuclear localization of TFs, on-the-fly nuclear detection and segmentation is followed by random nucleus selection within user-defined size and brightness thresholds. Centered overview and small FOV images of user-defined size (yellow square) are acquired in both densely and sparsely labeled POI channels. 2) An SMT illumination sequence is triggered in the small FOV. On-the-fly detection, localization and tracking of single molecules allows to reject too dense localizations (left; red box) and to determine illumination parameters yielding sparse and trackable single-molecule densities (right; green box). 3) Post-acquisition, nuclei in small FOVs of densely labeled epifluorescence POI images are resegmented and masks are propagated to all channels. Manual quality control (QC) is performed to exclude nuclei with erroneous segmentation masks or uncharacteristic texture, and bright autofluorescent debris (red), and to subject only correct nuclei to further analysis (green). Nuclear masks are used to extract cellular morphological characteristics (nuclear area, eccentricity) and mean TF intensities as a proxy for relative TF expression levels, and to filter out background single molecules outside of masks. 4) Cell level-based diffusion analysis provides single-cell diffusion spectra and diffusion parameters (fraction bound, modal diffusivity) and correlates them with cellular features extracted in 3). Scale bars: 50 μm (1); 5 μm (3). **(E)** Single-cell diffusion heatmaps for 8 combined automated SMT experiments for Hes1 (left; light turquoise; combined *n*=335 cells with 9,15,17,32,64,33,60,37,68 cells per experiment) and for 3 combined automated fast SMT experiments for Hnf4g (right; dark turquoise; combined *n*=186 cells with 68,18,100 cells per experiment). Cells are ordered according to increasing fraction bound from bottom to top. **(F)** Mean diffusion spectra for the combined experiments from E) for Hes1 (light turquoise) and Hnf4g (dark turquoise). Hes1: Bootstrap analysis of 8 combined experiments with *n*=335 cells for Hes1 and 3 combined experiments with *n*=186 cells for Hnf4g determined a mean fraction bound of 31.7% (95% CI: 26.3-37.3%) and 56.0% (95% CI: 52.4-58.9%), respectively.

To interrogate and compare the diffusive behavior of the two lineage TFs Hes1 and Hnf4g in a differentiating cell population, we generated stable organoid lines expressing the absorptive lineage TFs Hes1, promoting absorptive lineage commitment, or Hnf4g, providing enterocyte identity (Fig. 2B). To ensure that their overexpression does not induce obvious differentiation defects, we monitored the morphology of 3D organoids for two months after line establishment. They continued to display a budding morphology similar to wild-type (WT) organoids (Fig. S2A,B). In addition, we performed immunostaining in 2D EMCs to confirm the presence of ISCs, proliferative and secretory progenitor, secretory Paneth cells, and absorptive enterocytes. Similar to WT organoids, all investigated cell types were present in the stable Hes1 and Hnf4g lines, showing that no major cell lineage or cell type composition bias was introduced by the modest overexpression of absorptive lineage TFs (Fig. S2C).

Next, we subjected the Hes1 line to SMT. Here, a typical manual SMT experiment of 15 randomly chosen cells showed a large variation in the peak in diffusion coefficient (Fig. 2C; Fig. S3). Due to the great heterogeneity of cell types represented in 2D enteroids, the number of measured cells needed to be increased by a magnitude to provide accurate statistical sampling.

### Automated SMT samples hundreds of cells and extracts morphological features for a single-cell diffusion analysis in a heterogenous differentiation model

To achieve this, we developed an automated imaging and analysis pipeline for HILO-based SMT of 2D monolayer cultures on a Nikon TI microscope (Fig. 2D). We programmed the microscope to raster in a grid-like pattern over the sample and acquire overview images in the bulk TF channel, allowing the detection of nuclei. Nuclear segmentation using StarDist^58^ was followed by the automatic selection and centering of a nucleus in the field-of-view (FOV). Thresholds for nuclear size and brightness excluded bright autofluorescent debris or dead cells. A smaller region of interest (ROI) was acquired in both densely and sparsely labeled TF channels, followed by triggering of a user-defined SMT illumination sequence (Fig. 2D-1). On-the-fly localization density assessment served as quality control (QC) for the selection of trackable staining and SMT conditions (Fig. 2D-2), whereby tracking of single molecules was performed using quot^59^. Post-acquisition, nuclear segmentation masks served for a manual QC to exclude cells with erroneous segmentation masks or atypical nuclear textures, or that were out-of-focus. In addition, nuclear features, such as size and eccentricity, and the mean intensity-based relative TF expression level were determined (Fig. 2D-3). Finally, single-molecule diffusion analysis based on saSPT^59^ and a single-cell level correlation of morphological and diffusive parameters was performed (Fig. 2D-3).

**Figure 3:**
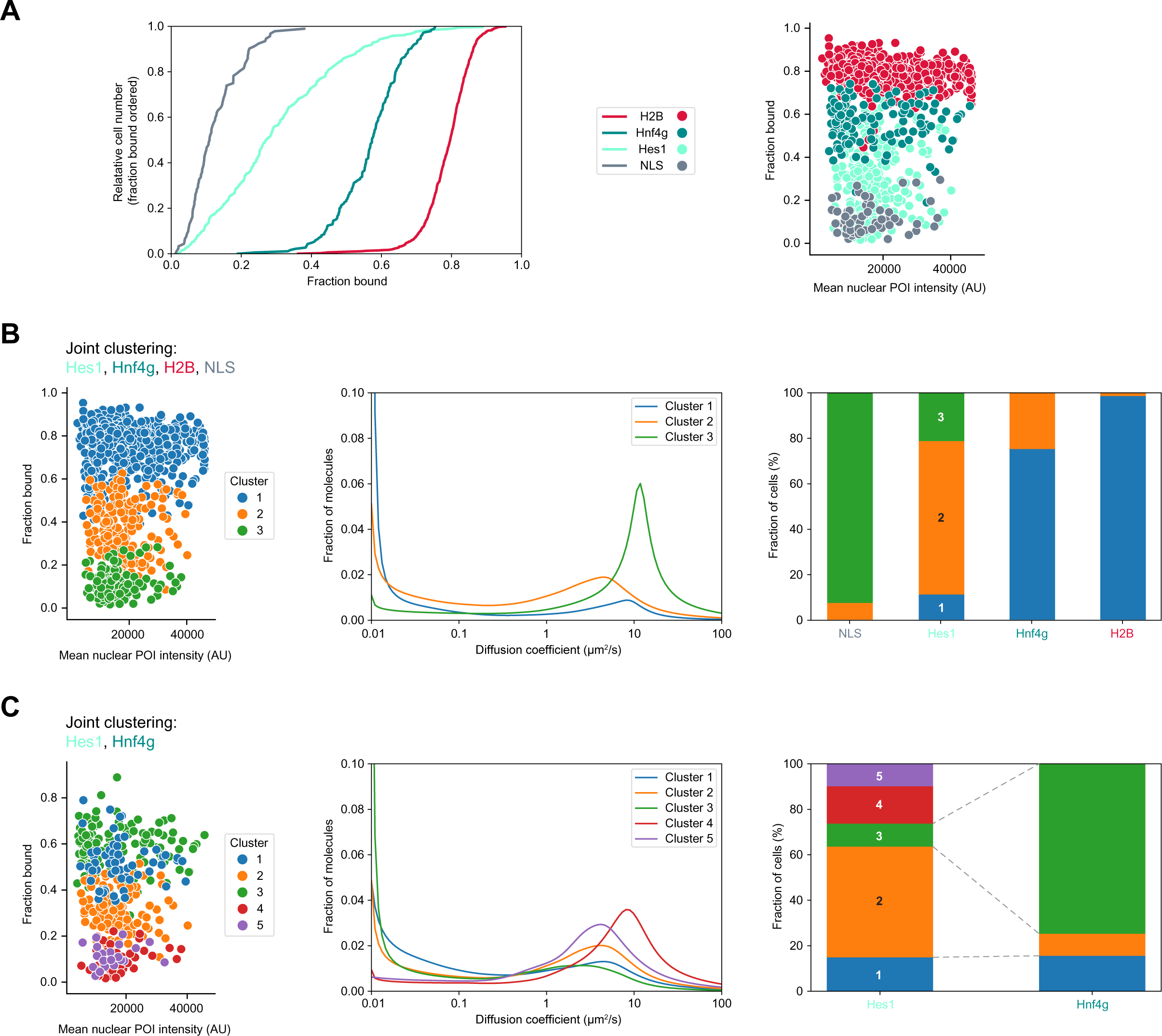
Cluster analysis of single-cell diffusion spectra reveals expression level-independent contrasting diffusive behaviors of Hes1-Halo and Hnf4g-Halo different from chromatin-bound H2B and freely diffusing NLS. **(A)** Comparative cell level-based distribution of fractions bound for Halo-NLS (gray), Hes1-Halo (light turquoise), Hnf4g-Halo (dark turquoise), and H2B-Halo (red) determined by SMT in 2D EMCs. Left: Fraction bound distributions are plotted in relative cell numbers with an increasing fraction bound from bottom to top. Right: Fractions bound for each cell are plotted against the mean nuclear POI intensity. **(B)** Joint hierarchical clustering of Hes1, Hnf4g, H2B, and NLS based on single-cell diffusion spectra using the Jensen-Shannon distance metric. Left: Fractions bound for each cell against the mean nuclear POI intensity with color-coded diffusion clusters. Middle: Mean diffusion spectra of each cluster. Right: Distribution of cells into diffusion clusters for each POI. Cluster: 1 (blue), 2 (orange), 3 (green). **(C)** Joint hierarchical clustering of Hes1 and Hnf4g as in (B). Left: Fractions bound of each cell against the mean nuclear POI intensity with color-coded diffusion clusters. Middle: Mean diffusion spectra of each cluster. Right: Distribution of cells into diffusion cluster for each POI. Cluster: 1 (blue), 2 (orange), 3 (green), 4 (red), 5 (purple). Shown are *n*=92 cells from 3 combined manual experiments for NLS (*n*=16,42,34 cells) (A,B), *n*=335 cells from 8 combined automated experiments for Hes1 (*n*=9,15,17,32,64,33,60,37,68 cells per experiment) (A-C), *n*=186 cells from 3 combined automated experiments for Hnf4g (*n*=68,18,100 cells per experiment) (A-C), and *n*=1022 cells from 5 combined automated experiments for H2B (*n*=134,90,92,355,351 cells per experiment) (A,B). Cells for NLS are from manual experiments in Fig. 1G and Fig. S1, whereas cells for Hes1, Hnf4g and H2B are from automated experiments in Fig. 2E,F and Fig. S5 (Hes1, Hnf4g), and Fig. S4 (H2B).

Using H2B as a test case, we demonstrated the power of this first completely academic-scale automated imaging and analysis platform for live-cell SMT^60,61^ by recording more than 7000 cells in 5 combined experiments. About 1000 of them passed QC and were selected for diffusion analysis, confirming our results from manual SMT (Fig. 1G), whereby about 60% of them were fully included in the ROI and subjected to correlative diffusion and cellular feature analysis (Fig. S4).

### Automated SMT reveals a contrasting diffusive behavior of the early and late absorptive TFs Hes1 and Hnf4g

Interestingly, by imaging over a magnitude more cells, we revealed differences in the diffusive behavior of Hes1 and Hnf4g in 2D EMCs: Hes1 displayed a large cell-to-cell variability with peaks in the diffusion spectra ranging from freely diffusing (10 μm^2^/s) to chromatin-bound (0.1 μm^2^/s), suggesting the presence of different diffusive Hes1 states across the differentiating population (Fig. 2E; Fig. S5A,B). In contrast, the diffusive behavior of Hnf4g was more uniform within most cells being characterized with a peak in diffusion coefficient in the immobile spectrum, indicating less variability across non-differentiated and differentiated cells (Fig. 2E; Fig. S5A,B). Further comparative assessment revealed a lower fraction bound of 31.7% for Hes1 and a higher fraction bound of 56.0% for Hnf4g (Fig. 2F; Fig. S5C), suggesting that on average a larger fraction of Hnf4g molecules is bound to DNA. Moreover, freely diffusing Hnf4g was characterized by a slower and broader diffusing peak than Hes1, indicating the presence of Hnf4g in slower moving protein complexes (Fig. 2F; Fig. S5A,B).

Despite having ruled out stark differentiation defects in our stable organoids (Fig. S2), we investigated whether the demonstrated diffusive TF behavior is also reflected upon short-term overexpression of TF transgenes. To this end, we generated stable organoid lines expressing the TFs under a doxycycline-inducible promoter (Fig. S6A). While a lower fraction bound for Hnf4g could be explained by protein steady states not having been reached 1d after induction, our previously observed differences in the population-averaged and cell level-based diffusive behavior of Hes1 and Hnf4g persisted (Fig. S6B-D).

### Cluster analysis of single-cell protein diffusion reveals characteristic expression level-independent diffusive behaviors for Hes1 and Hnf4g

As expected for TFs, the average bound fractions for Hes1 and Hnf4g lie in between those for H2B and NLS (Fig. 3A; Fig. S7A,B). Comparing their diffusive behavior on the single-cell level revealed that Hes1 stretches out to about more than three times the range of fractions bound determined for H2B, NLS, and Hnf4g (Fig. 3A). Importantly, these fraction-bound distributions were independent of the POI expression level, ruling out any potential effect due to heterogeneity stemming from the initially polyclonal organoid lines (Fig. 3A).

To further compare diffusive protein behavior, we hierarchically clustered the single-cell diffusion spectra using the Jensen-Shannon distance metric^62^. To validate our clustering method, we applied it to all four proteins together and observed three clusters (Fig. 3B; Fig. S7C) with H2B and NLS controls almost completely comprising the slowest (cluster 1) and fastest diffusing cluster (cluster 3; Fig. 3B), respectively. While about three quarters of Hnf4g cells clustered together with H2B, one fourth co-clustered with the majority of Hes1 with intermediate diffusivity (cluster 2). Consistent with its large spread in fractions bound (Fig. 3A), Hes1 was present in all three clusters, further underscoring its large variability in peak diffusion coefficients (Fig. 3B).

Zooming in to compare only Hes1 and Hnf4g, we clustered these two TFs together (Fig. 3C; Fig. S7D). While about three quarters of Hnf4g cells fell into one cluster (3) with the rest being split between clusters 1 and 2, Hes1 was represented in all five clusters with about half of Hes1 cells falling into cluster 2 (Fig. 3C). This further confirmed a more uniform diffusive behavior of Hnf4g in contrast to a larger cell-to-cell variability in Hes1 diffusion. Notably, despite similar fractions bound, the two fastest diffusing clusters, solely occupied by Hes1, were characterized by different peaks in the fast-diffusing spectrum (Fig. 3C), further demonstrating the power of our clustering approach to not only rely on the fraction bound but also on the shape of the diffusion spectrum. We speculate that these clusters could represent freely diffusing monomeric (cluster 4) versus dimeric (cluster 5) Hes1^29^.

### Cells with a higher fraction of bound Hes1 display nuclear characteristics of proliferative stem and early progenitor cells

The wide range in fractions bound for Hes1 raised the question whether a different diffusive behavior might reflect differentiation status. We therefore investigated a potential correlation between diffusive parameters (fraction bound, modal diffusivity) and cellular morphological features (nuclear area, eccentricity) at the single-cell level (Fig. 4A). Interestingly, we identified a subpopulation of Hes1 cells with higher fractions bound characterized by smaller nuclei or a larger eccentricity (Fig. 4A). In contrast, we detected a small subpopulation of Hnf4g with lower fractions bound characterized by smaller nuclei (Fig. 4B). Intrigued by these observations, we quantified the relationship between morphological parameters and the proliferation marker Ki67^63^ (Fig. 4C). This indeed revealed negative or positive correlations between mean nuclear Ki67 intensity and nuclear area or eccentricity, respectively (Fig. 4C), providing indirect evidence that cells with smaller nuclei and a higher eccentricity are stem or early progenitor cells residing in proliferative centers (Fig. 1A). For Hes1, this is indicative of a proliferative subpopulation characterized by a higher fraction bound, consistent with Hes1 acting early during absorptive differentiation to confer absorptive lineage commitment. In contrast, for Hnf4g, this suggests that larger fractions bound would be associated with non-proliferative differentiated cells, consistent with Hnf4g acting later during absorptive differentiation to confer enterocyte identity.

**Figure 4:**
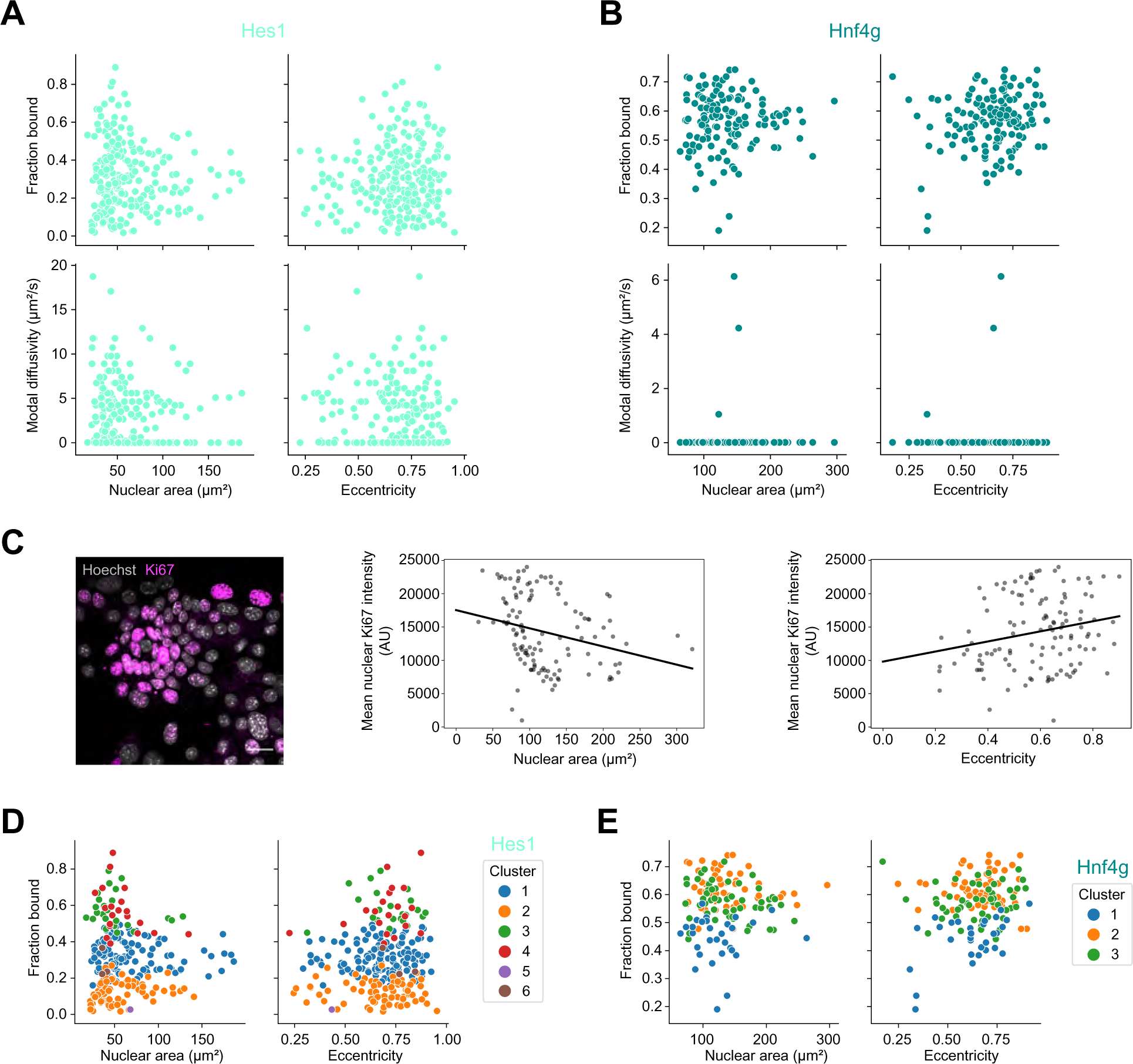
Cell level-based extraction of morphological and diffusion parameters reveals distinct subpopulations with a higher fraction bound of Hes1 with proliferative stem and early progenitor cell characteristics. (A,B) Single-cell level-based correlation of SMT-derived diffusion parameters (fraction bound, modal diffusivity) for Hes1 (A) or Hnf4g (B) with morphological characteristics (nuclear area, eccentricity) derived from nuclear segmentation masks (dense TF-Halo labeling) in 2D EMCs. **(C)** Left: Confocal image of a 2D WT EMC immunostained for Ki67 (magenta) and co-stained with Hoechst (gray). A 160.44 μm x 160.44 μm-large FOV of the full-size image is shown. Scale bar: 20 μm. Middle: Negative correlation between mean nuclear Ki67 intensity and nuclear area. Y-intercept: 17539.28; slope: -27.31. Right: Positive correlation between mean nuclear Ki67 intensity and nuclear eccentricity. Y-intercept: 9819.73; slope: 7563.12. Brightness was adjusted for display purposes (left) but quantifications (middle, right) were performed on the non-adjusted full-size raw image. **(D,E)** Hierarchical clustering of Hes1 (D) and Hnf4g (E) based on single-cell diffusion spectra using the Jensen-Shannon distance metric. Fractions bound plotted against cellular morphological characteristics with color-coded diffusion clusters. Data plotted in (A,D) correspond to 8 combined automated experiments for Hes1 with *n*=9,15,17,32,64,33,60,37,68 cells and data plotted in (B,E) correspond to 3 combined automated experiments for Hnf4g with *n*=68,18,100 cells shown in Fig. 2E,F and Fig. S5.

To investigate the diffusive uniformity of these subpopulations, we applied our hierarchical clustering approach to Hes1 (Fig. 4D; Fig. S8A) or Hnf4g alone (Fig. 4E; Fig. S8C). While four major clusters were revealed for Hes1 (Fig. 4D; Fig. S8A), Hnf4g diffusive behavior comprised three clusters (Fig. 4E; Fig. S8C), further confirming a higher degree of complexity in the diffusion behavior of Hes1 across the cell population. Importantly, the three clusters revealed for H2B displayed no differences in cellular morphological features (Fig. S9A,B), consistent with H2B diffusivity not changing upon differentiation. Similar, cells across the whole range of nuclear area and eccentricity were found in all three Hnf4g diffusion clusters (Fig. 4E; Fig. S8C,D). Strikingly, cells in the two slower diffusing Hes1 clusters with larger fractions bound (cluster 3 and 4) were almost exclusively characterized by smaller nuclei and a larger eccentricity (Fig. 4D; Fig. S8A,B), further suggesting the presence of subpopulations with distinct Hes1 diffusive behaviors displaying morphological characteristics of proliferative ISC and progenitor cells.

### Probing molecular proximity by PAPA-SMT reveals predominantly chromatin-bound self-associated Hes1 and Hnf4g

While the two absorptive lineage TFs Hes1 and Hnf4g display different diffusive behaviors indicative of their early and late action along the enterocyte differentiation trajectory, both TFs are known to form homodimers^29,35^. To probe for their self-association in EMCs,we double-labeled the TF-Halo population with two different sender and receiver fluorophores for an SMT assay combined with PAPA^47,55^ (Fig. 5A). Here, upon shelving JFX650-labeled TF-Halo using far red light, a series of violet (direct reactivation (DR)) and green (PAPA) light pulses highlight either all JFX650-labeled single molecules (similar to standard SMT) or only those that are in close vicinity to JFX549-labeled single molecules, namely primarily TF-Halo oligomers (Fig. 5B; Fig. S11; Fig. S12A,B). For both Hes1 and Hnf4g, we detected faster and slower diffusing self-associated populations (Fig. 5C; Fig. S12C), indicative of diffusing homodimers. Most notably, for both POIs the fraction bound was increased to about 10 percentage points in the PAPA *versus* the DR condition (35.8±3.7% *vs.* 24.8±1.9% for Hes1; 63.1±2.0% *vs.* 53.8±1.4% for Hnf4g) (Fig. 5C,D), suggesting that self-association of both lineage TFs is occurring in the immobile, DNA-bound state. These increased bound fractions are likely underestimated due to the undetectability of self-association with endogenous unlabeled TFs and background PAPA signal of non-interacting TFs (Fig. S11). In our assay, these results nevertheless provide further evidence that dimerized DNA-bound states confer TF functionality and demonstrate the power of SMT and PAPA to probe TF function.

**Figure 5:**
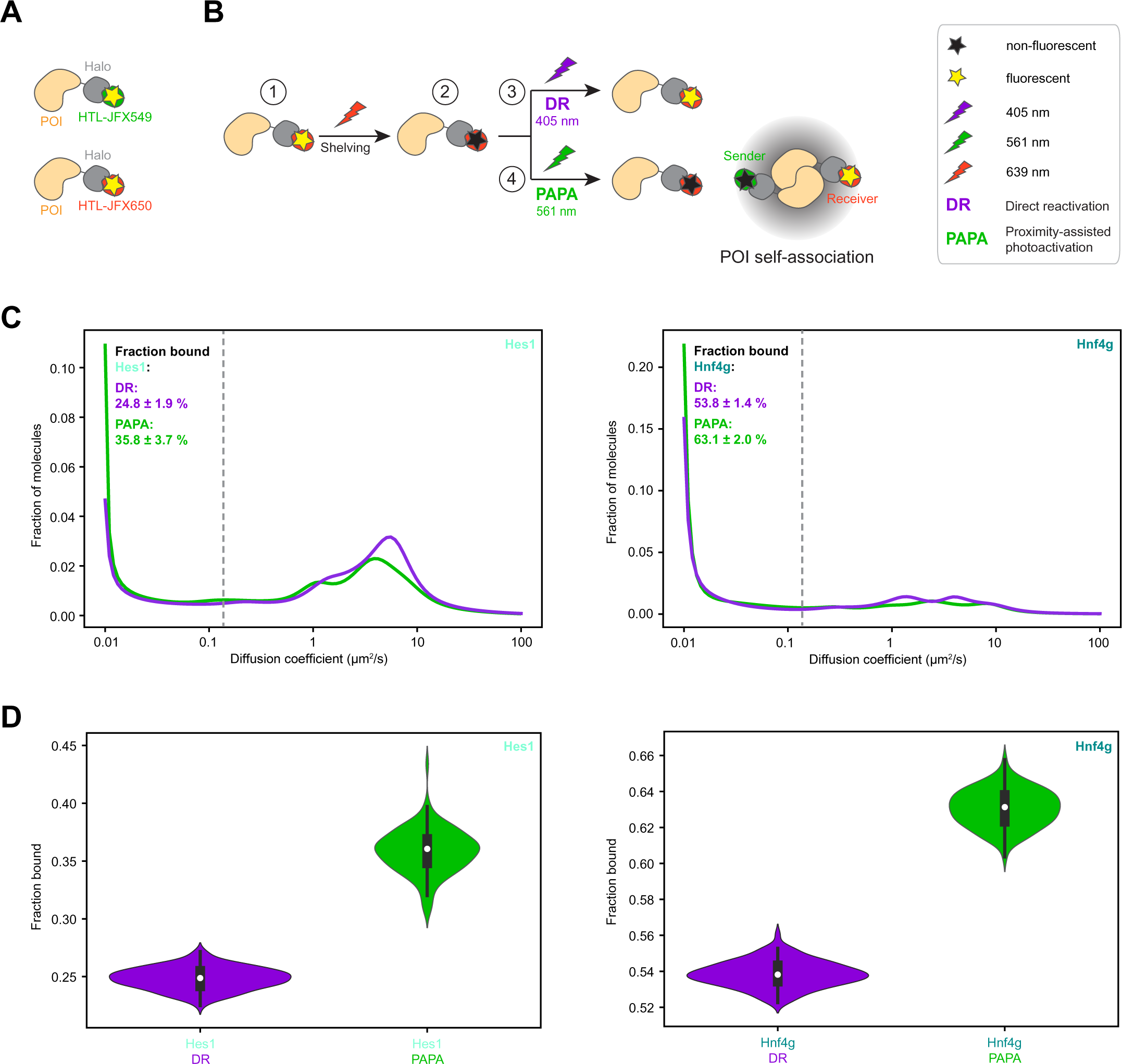
Proximity-assisted photoactivation (PAPA)-SMT reveals that self-associated Hes1 and Hnf4g are predominantly chromatin-bound. **(A)** Double labeling of POI-Halo (orange-gray) with two different HTL-coupled fluorophores (JFX549, sender, green; JFX650, receiver, red) for PAPA-SMT. **(B)** Principle underlying PAPA-SMT: 1) JFX650-labeled and far-red light emitting (yellow asterisk) receiver molecules (red) are put into the 2) dark state (shelving) through excitation (bleaching) with 639 nm light (red lightening arrow). 3) Direct reactivation (DR; purple) of dark JFX650-labeled receiver molecules with a violet 405 nm light pulse (purple lightening arrow) makes them detectable independent of any potential oligomerization states as in standard SMT. 4) PAPA illumination (green) of dark JFX650-labeled receiver molecules with a green 561 nm light pulse (green lightening arrow) makes them detectable only if in proximity (gray cloud) to a JFX549-labeled sender molecule (green). The most common though not exclusive scenario for proximity-based PAPA of the same POI is through self-association/oligomerization (see Fig. S11). **(C)** Mean PAPA (green) *versus* DR (purple) diffusion spectra for PAPA-SMT of double-labeled Hes1-Halo (left) or Hnf4g-Halo (right) in 2D EMCs. **(D)** Violin plots for fractions bound (median (white point); first/third quartile (whiskers)) of Hes1 (left) and Hnf4g (right) determined from DR (purple) *versus* PAPA (green) trajectories. Data in (C,D) are from 3 combined experiments with 232 cells for Hes1 with *n*=84,90,58 cells per experiment and 13471 DR and 3058 PAPA trajectories (fractions bound: DR: 25.0%, PAPA: 35.7%) and from 3 combined experiments with 326 cells for Hnf4g with *n*=60,131,135 cells per experiment and a total of 31024 DR and 12282 PAPA trajectories (fractions bound: DR: 53.7%, PAPA: 62.9%). Hes1: Subsampling of 3058 trajectories determined fractions bound to 24.9% (DR) and 35.7% (PAPA). Bootstrap resampling with 100 replicates determined fractions bound to 24.8±1.9% (DR) and 35.8±3.7% (PAPA); 2-sided *p*-value: 2.4e-7 (displayed in (C,D); left). Hnf4g: Subsampling of 12282 trajectories determined fractions bound to 53.3% (DR) and 62.9% (PAPA). Bootstrap resampling with 100 replicates determined fractions bound to 53.8±1.4% (DR) and 63.1±2.0% (PAPA); 2-sided *p*-value: 1.9e-13 (displayed in (C,D); right).

## Discussion

Using state-of-the-art fast SMT, we investigated the long-standing question of how the diffusive behavior of lineage TFs correlates with cell state transitions within multicellular tissue-resembling monolayer cultures recapitulating differentiation. Focusing on ISC differentiation during adult tissue homeostasis in the intestinal epithelium, we established live-cell HILO-based SMT to measure TF diffusion dynamics and PAPA-SMT to investigate their self-interaction in 2D EMCs derived from 3D mSIOs (Fig. 1; Fig. 2; Fig. 5). As resolving distinct diffusive subpopulations of cells in such heterogenous samples consisting of ISCs and differentiated cell types required sampling hundreds of cells (Fig. 2C), we developed an automated pipeline for SMT imaging and analysis (Fig. 2D).

While industrial-scale automated SMT for drug screening has recently been achieved^60,61^, its implementation in academia remained challenging due to limitations in budget and resources. Here, we overcame these obstacles by developing an automated imaging and analysis pipeline for HILO-based SMT on a standard inverted TIRF microscope compatible with stroboscopic illumination and streamlining our existing analysis packages (quot, saSPT)^59^. Cellular feature extraction (Fig. 2D) together with hierarchical clustering of protein diffusion spectra (Fig. 3B,C) allowed us to analyze single-cell TF diffusion behavior in relation to TF expression level (Fig. 3B,C) and cellular morphological characteristics distinct for stem/progenitor and differentiated cell types (Fig. 4; Fig. S10).

Applying our developed SMT pipeline to study the lineage-determining TFs Hes1 and Hnf4g, acting early and late during absorptive intestinal differentiation, respectively (Fig. 6A), we discovered opposing diffusive behaviors. The diffusion dynamics of Hes1, the key TF determining absorptive lineage commitment, were characterized by a large cell-to-cell variability (Fig. 2E; Fig. 3). Importantly, we resolved subpopulations with distinct Hes1 diffusion and cell morphology characteristics independent of its expression level. Consistent with its early absorptive lineage-determining role, stem/progenitor cells were characterized by a higher fraction bound of Hes1 molecules in contrast to a lower fraction bound in differentiated cells (Fig. 4D). Despite the absence of distinct separately clustering intermediate populations, these findings suggest that the Hes1 diffusion dynamics can be indicative of the progression of differentiation to enterocytes with a decrease in fraction bound with gained maturity. In addition, the overall lower fraction of DNA-bound Hes1 molecules of ∼32% (Fig. 2E; Fig. 3) is consistent with the requirement for early lineage decisions being reversible to enable dedifferentiation upon damage to the stem cell compartment^64^ and in line with ISCs having been characterized as broadly permissive epigenetic cell states, enabling fast transitions to differentiated states^36,37^. In contrast to Hes1, the more uniform dynamics of the enterocyte determinant Hnf4g across the cell population with an overall larger fraction bound of ∼56% (Fig. 2F; Fig. 3) seem to support later stages of the differentiation of absorptive progenitors to enterocytes, which need to be irreversible to lock in enterocyte identity and ensure their key role in nutrient absorption in the intestine (Fig. 6B). Differentiation of ISCs to enterocytes is considered as the shortest path in intestinal differentiation space^65^ with early absorptive lineage commitment already happening in crypts^66^. Nevertheless, major changes in the gene expression pattern need to occur to achieve enterocyte differentiation, likely induced by the action of a single or few TFs^18^, whereby Hnf4g was found to ensure long-range enhancer-promoter interactions^67^.

**Figure 6:**
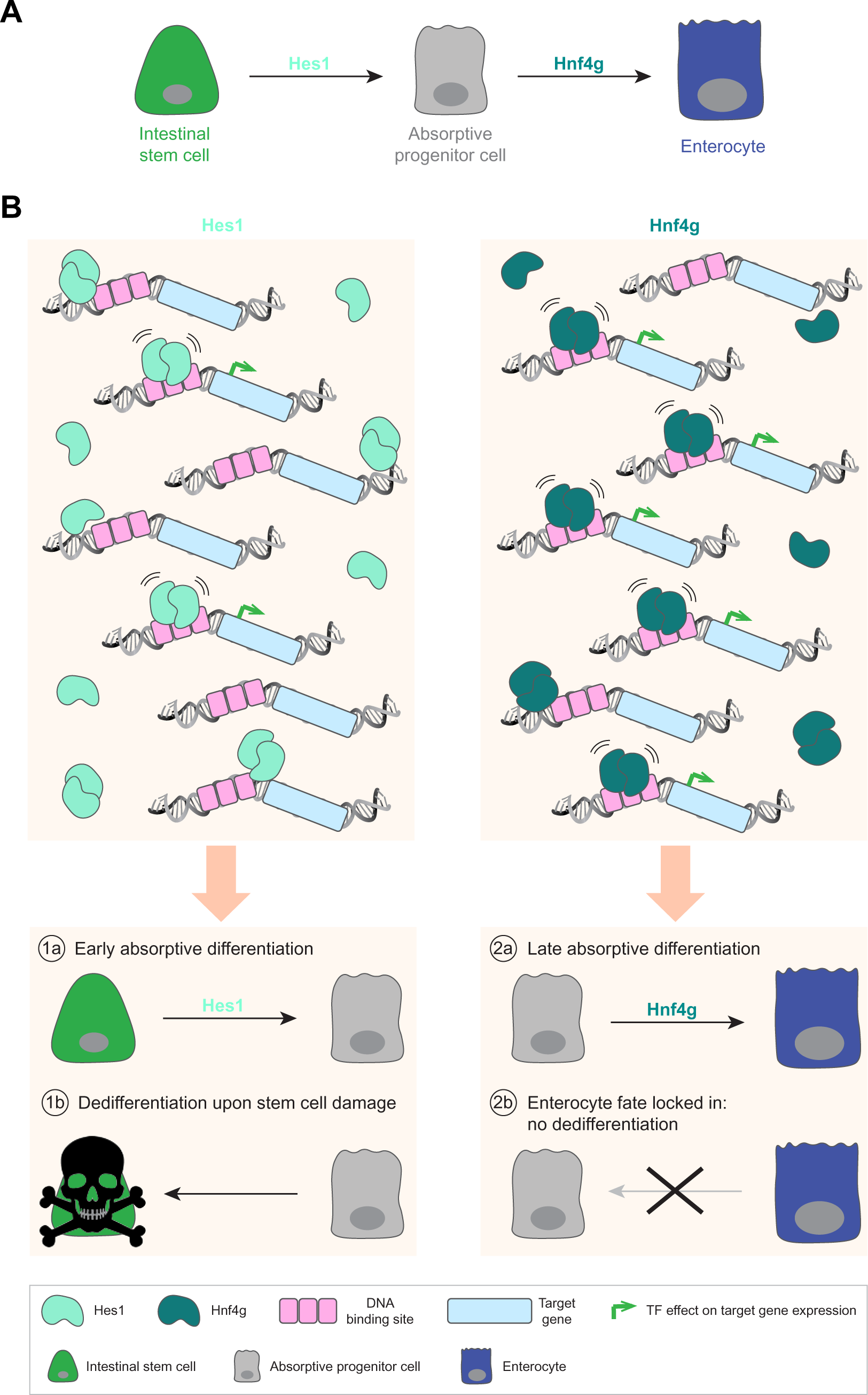
Summary and model of how the contrasting diffusive behavior of Hes1 and Hnf4g could support early *versus* late absorptive differentiation in the small intestine. **(A)** Hes1 (light turquoise) is the key absorptive TF in the mouse small intestinal epithelium regulating lineage commitment of ISCs (green) to absorptive progenitor cells (gray). Hnf4g (dark turquoise) acts later along the absorptive differentiation trajectory by conferring enterocyte identity (blue). **(B)** Top: Large cell-to-cell variability in the diffusive behavior of Hes1 with an overall lower fraction of DNA-bound molecules. The diffusive behavior of Hnf4g is more uniform and characterized by a higher fraction bound. For both TFs, the immobile, DNA-bound, and hence transcriptionally active moiety are homodimers. Bottom: SMT data-driven model for the opposing diffusive behavior of Hes1 and Hnf4g supporting gene regulatory requirements for early, plastic *versus* late, irreversible absorptive differentiation to enterocytes: The overall lower fraction bound of Hes1 supports its function during early absorptive differentiation (1a) and enables dedifferentiation of progenitor cells to ISCs upon damage to the stem cell compartment (1b). In contrast, the overall higher fraction bound of Hnf4g supports late absorptive differentiation to enterocytes (2a) and irreversibly determines enterocyte fate (2b).

Taken together, our results suggest different modes of gene regulation through the absorptive lineage-determining TF Hes1 or enterocyte identity-conferring TF Hnf4g during early and late absorptive differentiation towards enterocytes (Fig. 6A). We thus propose a model in which this differential diffusion behavior supports early, plastic, *versus* late, irreversible cell fate decisions (Fig. 6B). Despite the association of aberrant TF expression with developmental disorders and tumorigenesis^3,4^, our results suggest that lineage TF expression alone is not a sufficient determinant of differentiation. Instead, varying TF diffusive behavior confers differentiation and can modulate early *versus* late differentiation within the same lineage.

While Hnf4g functions only in the gastrointestinal tract, it will be interesting to investigate whether the revealed principle of Hes1 dynamics holds true in other differentiating tissues^68–70^. Independently, differential TF dynamics have been shown in various embryonic and adult differentiation contexts^44,49,71^, suggesting a broader applicability for TF dynamics as a regulatory principle during differentiation. Arguing against simple gene expression regulation through binding of a *trans*-acting TF to *cis*-regulatory elements, exciting future avenues will include interrogating the molecular mechanisms underlying these differential TF dynamics, such as post-translational modifications and association with co-factors^2^, or the formation of protein microenvironments^72^, as well as to quantitatively probe the lineage TF concentration dependence^73^ for cell state transitions during differentiation and malignant transformation.

Dimerization of Hes1 is required for DNA binding^29^. However, direct evidence for Hes1 dimer formation in the intestine has been lacking. Investigations of protein oligomerization have so far only been possible by biochemical crosslinking and fractionation in fixed and lysed populations of thousands of cells or by live-cell single-molecule FRET (Förster resonance energy transfer) operating only at very short distances^74^. Here, we performed the first live-cell protein self-association study of a lineage TF in a differentiation context and demonstrated that self-associated Hes1 is indeed enriched in the chromatin-bound fraction (Fig. 5C,D), confirming Hes1 dimers as the functional unit in intestinal gene expression regulation. Dimerization inhibitors could further clarify the role of Hes1 dimersin transcription regulation^75^. Further developing PAPA-SMT towards using orthogonal tags for different labeling of two putative interaction partners^47,55^ will make it possible to perform biochemistry in live cells and interrogate complex protein interaction networks within tissue-recapitulating systems, such as a potential cell type-dependent homo- (Fig. 5C,D) and heterodimer formation of Hnf4g and its paralog Hnf4a with reportedly redundant yet still controversial roles during intestinal differentiation^19,76^.

Overall, we demonstrated that high-throughput fast SMT in 2D EMCs represents a powerful gateway into dissecting molecular mechanisms within a multicellular tissue-resembling context (Fig. 7A). In combination with small-molecule or genetic perturbations, our approach could be suitable for preselecting promising protein candidates in functional assays prior to further exploiting their dynamics in the 3D tissue context. We anticipate studies of (de-)differentiation during intestinal epithelial homeostasis and regeneration. Furthermore, we envision a broad application potential of our automated SMT pipeline to other 2D differentiation systems, such as 2D human intestinal differentiation models^77^ for studying healthy *versus* diseased conditions and drug screening, monolayer cultures recapitulating differentiation of other tissues^78,79^, or 2D micropatterned gastruloids^80–82^ to investigate TF dynamics during embryonic patterning (Fig. 7B).

**Figure 7:**
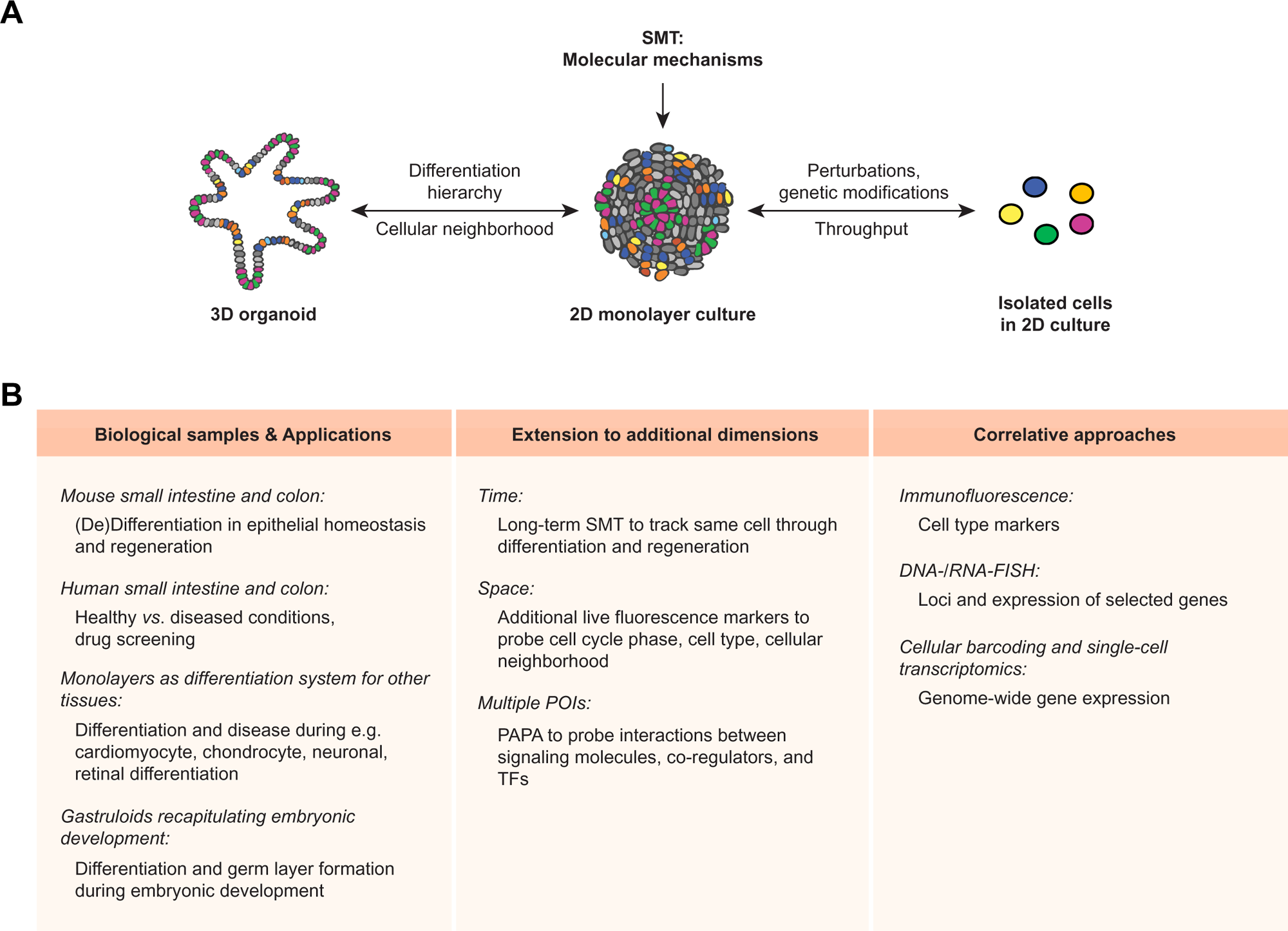
Methodological summary and outlook to future applications and developments. **(A)** Summary: Our broadly applicable automated live-cell SMT framework enables the study of molecular mechanisms by measuring single-molecule dynamics within *in vitro* 2D tissue models at the single-cell level. **(B)** Outlook: Future applications and developments.

Future developments will extend our experimental framework to additional time and space dimensions. We envision establishing long-term SMT to allow measurements at multiple time points by tracking cells through differentiation. Additional live fluorescence markers will help probing TF dynamics in relation to cell type or cell cycle, largely increasing the gain in context information. Finally, the development of correlative approaches coupling SMT measurements with complementary immunofluorescence, DNA- or RNA-FISH, or single-cell transcriptomics methods has the potential of revolutionizing tissue biology by bridging scales down to the single-molecule level (Fig. 7B).

In conclusion, the automated SMT pipeline developed in this study has allowed us for the first time to determine diffusion properties of lineage TFs in a heterogenous differentiating tissue-resembling model system, to correlate them with TF expression level and cellular morphological characteristics, and to reveal novel principles of lineage TF dynamics in modulating stem cell differentiation along the same differentiation trajectory. Our approach bridges scales between single molecules and simplified *in vitro* differentiation-recapitulating monolayer cultures. Together with its versatility and compatibility with high-throughput assays and perturbations, we envision a broad application potential of our technology to study the molecular mechanisms of fundamentally important processes of life, such as tissue homeostasis and regeneration upon injury, embryonic development, and malignant transformation.

### Limitations

While 2D EMCs are composed of all epithelial cell types of the small intestine and recapitulate important features of the gut, they represent a simplistic *in vitro* tissue model with reduced dimensionality. Hence, validation experiments in 3D organoids or tissues are recommended. This requires different microscopy techniques capable of imaging single molecules inside thicker specimen, as HILO-based SMT is limited to the first cell layer above the cover slip.

Our SMT approach relies on POI-Halo fusions. While other self-labeling tags, such as SNAPf^83^ or photoconvertible fluorescent proteins^84^, could be used, they all require POI fusion proteins and testing for functionality. Here, we used slight overexpression of transgene-encoded POI-Halo in addition to unlabeled endogenous POIs. Thus, the presence of a mixed protein population needs to be considered during data interpretation, and validation experiments are required, as we performed in this study (Fig. S2; Fig. S6B-D). Further improvements could rely on endogenously tagging the POI^85^ to study its molecular dynamics at physiological expression levels.

## Material and Methods

### Resource availability

#### Lead contact

Further information and requests for resources and reagents should be directed to and will be fulfilled by the lead contact.

#### Materials availability

Plasmids generated in this study are available upon request with a completed material transfer agreement and with reasonable compensation by the requestor for shipping. There are restrictions to the availability of stable organoid lines generated in this study due to the lack of an external centralized repository for its distribution and our need to maintain the stock.

### Experimental model and study participation details

#### Mice

Female and male C57BL/6J mice were obtained from The Jackson Laboratory (strain #000659, RRID:IMSR_JAX:000664), housed and bred in an AAALAS-certified level 3 facility on a 14 h light cycle. Pups were weaned 21 d after birth and housed with four littermates per cage. Male offspring was used for mouse small intestinal crypt isolation and organoid generation at an age of 8-16 weeks. All procedures to maintain and use the mice were approved by the Institutional Animal Care and Use Committee of the University of California, Berkeley (IACUC protocol number AUP-2015-09-7988-2).

#### Cell lines

L-Wnt3a cells (CRL-2647, ATCC) were used to produce Wnt3a-conditioned medium. Hek293T Lenti-X cells (632180, Takara) were used for making lentivirus. Cell lines were obtained via the UC Berkeley Cell Culture Facility. Cell lines and organoids were confirmed to be mycoplasma-free by regular PCR testing.

### Method details

#### Cloning of DNA constructs

For the production of lentivirus to deliver POI-Halo transgenes into organoids, the third generation lentiviral pHAGE vector originally developed in the lab of Richard Mulligan^86^ was used together with the second generation lentiviral packaging plasmid psPAX2 (gift from Didier Trono (Addgene plasmid # 12260; http://n2t.net/addgene:12260; RRID:Addgene_12260)) and the VSV-G envelope expressing plasmid pMD2.G (gift from Didier Trono (Addgene plasmid # 12259; http://n2t.net/addgene:12259; RRID:Addgene_12259)). For constitutive expression of POI-Halo-V5, a pHAGE L30 IRES Puro backbone was created and POI-Halo-V5 transgenes (H2B-Halo-V5, V5-Halo-NLS, Hes1-Halo-V5, Hnf4g-Halo-V5) were cloned into this vector to be expressed under the weak L30 promoter. For Hes1, the DNA sequence encoding the reference 282 aa isoform of UniProt entry P35428 was cloned (NCBI Gene ID 15205, NM_008235.3 isoform 1). For Hnf4g, the DNA sequence encoding the reference 408aa isoform of UniProt entry Q9WUU6 was cloned (NCBI Gene ID 30942, NM_013920.3 isoform 2). For doxycycline-inducible expression of POI-Halo-V5 constructs, a pHAGE TetOnDual backbone was created and POI-Halo-V5 transgenes (Hes1-Halo-V5, Hnf4g-Halo-V5) were cloned into this vector. The TetOnDual backbone was based conceptually on the bidirectional expression vector XLone^87^ but modified to contain the TRE3GV Tet responsive promoter to drive doxycycline-inducible transgene expression in one direction and the PGK TetOn 3G P2A Puro cassette to drive constitutive expression of the TetOn 3G transactivator as well as the puromycin resistance gene in the other direction. The final constructs were confirmed by Sanger sequencing. For further details, the plasmid maps including the cloning history will be made publicly available upon acceptance of this manuscript for publication in a peer-reviewed journal.

#### Lentivirus preparation

For lentivirus generation, low-passage 3.5×10^6^ Hek293T Lenti-X cells (632180, Takara) were seeded per 10 cm dish (Corning, cat.# 08-772E) in high-glucose Dulbecco’s modified eagle medium (DMEM) containing GlutaMAX-I (Gibco, cat.# 2537044), supplemented with 10% (vol/vol) fetal bovine serum (FBS; HyClone, cat.# SH30910.03, LOT# AXJ47554), and 1 mM sodium pyruvate (Gibco, cat.# 11360070) for transfection on the following day. The medium was changed 2 h prior to transfection. Lentivirus packaging and production was based on 2^nd^ generation lentivirus plasmids (see ‘Cloning of DNA constructs’). For calcium phosphate transfection, a plasmid cocktail consisting of 10 μg transfer vector containing the transgene in the pHAGE backbone and 3.5 μg and 6.5 μg of the two lentiviral packaging plasmids pMD2.G and psPAX2, respectively, was prepared and added to 426.5 μL 2:1 (vol:vol) 0.1x Tris (Fisher Scientific, cat.# BP152-5)-ethylenediaminetetraacetic acid (EDTA; Fisher Scientific, cat.# S311-500) (TE): ddH_2_O in a 15 mL conical tube. 50 μL 2.5 M CaCl_2_ (Millipore Sigma, cat.# C4901) were added to the mix, followed by quickly vortexing. 500 μL 2x Hepes buffered saline (HBS; pH 7.12; Millipore Sigma, cat.# 51558) were added dropwise to the mix while vortexing. The mixture was let sit for 15 min at room temperature (RT). Upon resuspending with a 1 mL pipette, the transfection mixture was added dropwise to the cells and the plate was moved horizontally and vertically for mixing. Cells were incubated at 37°C and 5% CO_2_ in a cell culture incubator for 2 d. Supernatants (SNs) containing secreted lentivirus were collected and filtered through a syringe filter with 0.45 μm diameter (Nalgene, cat.# 09-740-106) into a 50 mL conical tube. To concentrate the lentivirus preparation, 3.3 mL of Lenti-X concentrator (Takara, cat.# 631231) were added to 10 mL of filtered SN. Gentle mixing was achieved by inverting the tube, followed by incubation at 4°C over night (ON). The concentrated SN was centrifuged at 4°C for 45 min at 4000 g. Upon removal of the SN, pellets were kept on ice, resuspended in 100 μL phosphate buffered saline (PBS) without Ca^2+^ and Mg^2+^ (D-PBS; Stem Cell Technologies, cat.# 37350) and 20 μL aliquots were prepared and stored at -80°C.

#### Mouse small intestinal crypt preparation for growing small intestinal organoids

Male mice at an age of 8-16 weeks were euthanized and placed on ice. Upon opening their abdomen, the small intestine was dissected (leaving out the most proximal and distal 4 cm). The intestine was placed in a dish containing pre-cooled D-PBS supplemented with 1% (vol/vol) Penicillin/Streptomycin (Pen/Strep; Gibco, cat.# 15070063), and fat and mesenteric tissue were removed. Using a 18 Gauge gavage needle (Fine Science Tools, cat.# 1806150), the intestine was cleared by flushing with D-PBS containing Pen/Strep.

Upon placing the intestine in another dish containing fresh pre-cooled D-PBS with Pen/Strep, it was cut open longitudinally and laid open, followed by further washes and removal of large villi through moving the intestine across the dish. After repeating this procedure three times, the intestine was cut into 2-4 mm long pieces which were collected in 15 mL pre-cooled D-PBS containing Pen/Strep in a 50 mL conical tube. Using a 10 mL pipette pre-wetted in 1% (vol/vol) bovine serum albumin (BSA; Sigma, cat.# A0336-50mL) in D-PBS, the intestinal pieces were washed by pipetting up and down three times. Upon settling by gravity, the SN was carefully removed and replaced with 15 mL cold D-PBS. This procedure was repeated 15-20 times until the SN was clear. After removal of the SN, 25 mL Gentle Cell Dissociation Reagent (Stem Cell Technologies, cat.# 100-0485) were added, followed by incubation for 15 min on a rocking platform at 30 g. Upon settling by gravity, the SN was removed, and intestinal pieces were resuspended in 10 mL pre-cooled D-PBS containing 0.1% (vol/vol) BSA by pipetting up and down three times. After settling by gravity, the SN was removed, filtered through a cell strainer with a mesh size of 70 μm (Corning, cat.# 08-771-2) and collected in a 50 mL conical tube as fraction 1 and placed on ice. The procedure was repeated until fraction 4 was collected. The four fractions were inspected for the presence of crypts under a light microscope and those fractions enriched in crypts but not villi were centrifuged at 290 g and 4°C for 5 min. After removal of the SN, pellets were resuspended in 10 mL pre-cooled D-PBS containing 0.1% (vol/vol) BSA, transferred to a 15 mL conical tube, and centrifuged at 200 g and 4°C for 5 min. After removal of the SN, pelleted crypts were resuspended in 10 mL pre-cooled basal medium containing 0.1% (vol/vol) BSA. 1 mL of each resuspended fraction was used for counting crypt numbers and crypts were aliquoted to 500 crypts/mL, 1500 crypts/mL, and 3000 crypts/mL per fraction. Upon centrifugation at 200 g and 4°C for 5 min, the SN was removed and pelleted crypts were placed on ice. Each pellet was resuspended in 300 μL 90% (vol/vol) Matrigel Matrix for Organoid Culture (Corning, cat.# 08774406) in IntestiCult Organoid Growth Medium (Mouse) (Stem Cell Technologies, cat.# 06005) supplemented with 1% (vol/vol) Pen/Strep, and 50 μL were seeded per well of a 24-well plate (Corning, cat.# 353047). Upon incubation for 10 min at 37°C in a cell culture incubator, 650 μL IntestiCult supplemented with 1% (vol/vol) Pen/Strep were added per well. Culturing crypts for 5-10 d resulted in the formation of mSIOs.

#### Mouse small intestinal organoid culture

3D mSIOs were cultured in 24-well plates in 50 μL 90% (vol/vol) Matrigel Matrix for Organoid Culture in IntestiCult supplemented with 1% (vol/vol) Pen/Strep. mSIOs were passaged every week with one to two media changes in between depending on mSIO density. To this end, the medium was removed and replaced with 1 mL Gentle Cell Dissociation Reagent. Using a 1 mL tip pre-wetted with 1% (vol/vol) BSA in D-PBS, the Matrigel dome was physically broken apart and transferred to a 15 mL conical tube in which the content from all wells of the same condition were collected. Each well was washed with an additional 1 mL of Gentle Cell Dissociation Reagent which was then transferred to the collection tube. Collection tubes were incubated horizontally for 10 min at RT on a shaker set to 30 g. Following centrifugation at 290 g and 4°C for 5 min, tubes were placed on ice, the SN including the Matrigel layer above the organoid pellet was carefully removed, and the organoid pellet was resuspended in 5-10 mL pre-cooled basal DMEM/F-12 medium containing 15 mM HEPES (Stem Cell Technologies, cat.# 36254) using a 10 mL pipette pre-wetted with 1% (vol/vol) BSA in basal medium. Following another centrifugation at 200 g and 4°C for 5 min, tubes were placed on ice and the SN was carefully removed. Organoid pellets were resuspended in 90% (vol/vol) Matrigel Matrix for Organoid Culture in IntestiCult supplemented with 1% (vol/vol) Pen/Strep in a *n* times 50 μL volume with *n* being the number of wells of a 24-well plate to be seeded. 50 μL of organoid-Matrigel-medium suspension was seeded to the center of each well of a pre-warmed 24-well plate. Following incubation for 10 min at 37°C in the cell culture incubator, 650 μL of IntestiCult were added to each well.

#### Preparation of Wnt3a-conditioned medium

L-Wnt3a cells (CRL-2647, ATCC) were cultured in Advanced DMEM/F-12 (ADMEM; Gibco, cat.# 12634028) supplemented with 10% (vol/vol) FBS, 1% (vol/vol) GlutaMAX (Gibco, cat.# 35050061), 10 mM HEPES (Sigma-Aldrich, cat.# H0887-20mL), 1 mM N-acetyl-L-cysteine (Sigma-Aldrich, cat.# A9165-5G; 500 mM stock in ddH_2_O, sterile-filtered), and containing 500 ng/mL G418 (Gibco, cat.# 11811-031). For the preparation of Wnt3a-conditioned medium, cells were passaged 1:10 onto 10x 10cm dishes. Upon reaching 100% confluency, the medium was changed to 10 mL of the above medium without selection antibiotics. On d4 and d8 after medium change, the SN was harvested in 50 mL conical tubes and stored at 4°C. Media from both harvests were combined, sterile-filtered (sterile vacuum filter units, hydrophilic PVDF, 0.22μm, 500 mL; EMD Millipore, cat.# S2GVU05RE), aliquoted and stored at -80°C.

#### Lentiviral transduction of mouse small intestinal organoids and selection of stable organoid lines

To prepare mSIOs for lentiviral transduction, they were seeded in pre-transduction medium consisting of 50% (vol/vol) IntestiCult without Pen/Strep and 50% (vol/vol) Wnt3a-conditioned medium, supplemented with 100 mM nicotinamide (Sigma, cat.# N0636-100G; 1M stock in ddH_2_O), 10 μM ROCK inhibitor Y-27632 (Stem Cell Technologies, cat.# 72304; 10 mM stock in ddH_2_O), and 2.5 μM CHIR99021 (Stem Cell Technologies, cat.# 72052; 10 mM stock in dimethylsulfoxide (DMSO; Sigma-Aldrich, cat.# D2650)). Typically, 4 wells from a 24-well plate of 3D mSIOs were prepared per transduction condition. 4-5 d post-seeding, the medium was removed and embedded mSIOs were washed with D-PBS. After D-PBS removal, 500 μL TrypLE Express (Thermo Scientific, cat.# 12604013) were added to each well and Matrigel domes were dislodged with a 1 mL pipette. After incubation for 10 min at 37°C in the cell culture incubator, digested Matrigel domes were further broken down physically by vigorously pipetting first with a 1 mL pipette and then with a 200 μL pipette. Clusters were transferred to a 15 mL conical tube containing 10 mL ADMEM plus 5% (vol/vol) FBS for quenching. Following centrifugation at 1000 g and 4°C for 5 min, the pelleted cell cluster was resuspended in 230 μL transduction medium (pre-transduction medium containing 10 μg/mL polybrene (Millipore Sigma, cat.# TR-1003-G)) and transferred to one well of a pre-warmed 48-well deep well plate (Corning, cat.# 08-772-3D). 20 μL of lentivirus thawn on ice were added per well/transduction condition, followed by mixing by pipetting. The plate was sealed with parafilm and centrifuged for 1 h at 37°C and 600 g. Upon unsealing the plate, it was incubated for 6 h at 37°C in a cell culture incubator. Transduced organoids were pipetted up and down and transferred to a 1.5 mL tube containing 750 μL of basal DMEM/F-12 medium with 15 mM HEPES. Tubes were spun at 1000 g for 5 min at RT and placed on ice. After carefully removing the SN, transduced organoids were resuspended in 300 μL 90% (vol/vol) Matrigel Matrix for Organoid Culture in IntestiCult Organoid Growth Medium supplemented with 1% (vol/vol) Pen/Strep and seeded to six wells of a 24-well plate with 50 μL per well. Upon solidification of Matrigel domes through incubation for 10 min at 37°C in the cell culture incubator, domes were overlaid with 650 μL pre-transduction medium. 2-3 d post-transduction, the medium was exchanged to pre-transduction medium containing selection antibiotics (2 μg/mL puromycin (Thermo Scientific, cat.# A11138-03)). Medium was exchanged every 2-3 d until large, selected spheroids were obtained. Upon passaging selected spheroids, they were cultured in IntestiCult supplemented with 10 μM ROCK inhibitor Y-27632 and 2.5 μM CHIR99021 including selection antibiotics. After 2-3 d, the medium was exchanged to IntestiCult supplemented with 2.5 μM CHIR including selection antibiotics. After another 2-3 d, medium was exchanged to IntestiCult including selection antibiotics and thereafter used as culture medium of selected stable organoid lines.

Upon establishment of stable lines, mSIOs were cryopreserved in CryoStor CS10 (Stem Cell Technologies, cat.# 07931) and stored in liquid nitrogen. To this end, mSIOs were washed with D-PBS, Matrigel domes were broken apart physically by pipetting, and collected in a 15 mL tube placed on ice. Following centrifugation at 290 g and 4°C for 5 min, the SN was removed, and organoid pellets were washed in 5-10 mL of basal medium and centrifuged at 200 g and 4°C for 5 min. After removal of the SN, the organoid pellet was resuspended in CryoStor CS10, distributed to cryovials (150-200 organoids per 1 mL freezing medium to be thawn on 3-4 wells of a 24-well plate), and slowly frozen to -80°C before transfer to the liquid nitrogen on the following day.

For thawing organoids, the content of a cryovial thawn in a 37°C water bath was transferred to 5 mL basal medium containing 1% (vol/vol) BSA, centrifuged at 200 g for 5 min, and plated as described above.

Organoids were used for experiments until 8 weeks after line establishment or thawing of an early passage cryopreserved organoid aliquot.

#### Confocal imaging of POI-Halo organoid lines and TetON POI-Halo organoid lines

For confocal imaging of POI-Halo organoid lines, POI-Halo organoids were seeded in 10-20 μL 90% Matrigel droplets into 8-well Labteks II #1.5 (Nunc, cat.# 12-565-338) and overlaid with 500 μL IntestiCult. The medium was carefully exchanged after 3 d. On d6 post-seeding, organoids were stained with 500 nM HTL-JF635 (kind gift from Luke Lavis) for at least 1 h in DMEM/F-12 without phenol red (Gibco, cat.# 11039-021) and imaged 1-6 h after staining. No washout of the dye was required, as JF635 is fluorogenic, reducing luminal background fluorescence. For confocal imaging of TetON POI-Halo organoid lines, medium was exchanged to 200 μg/mL doxycycline (Sigma, D9891-1G) in IntestiCult on d4 post-seeding. Upon doxycycline-induced transgene expression for 1 d, TetON POI-Halo staining was performed on d5 post-seeding as described above with the exception that the staining/imaging medium contained 200 μg/mL doxycycline.

Confocal imaging of 3D mSIOs was performed on a Yokogawa CSU-W1 SoRa spinning disk with a Nikon Ti2 inverted microscope operated by the NIS Elements AR 5.42.03 software (Nikon). The microscope was equipped with a temperature- and CO_2_-controlled incubation chamber (Okolab), and the temperature was set to 37°C and the CO_2_ to 5% for imaging of live organoids. For BF organoid overviews (Fig. S2A,B), images were acquired using a Plan Apo λ 10× air objective (N.A. 0.45; Nikon). Multiple *z* planes (*xy* pixel size: 0.65 × 0.65 μm; *z* interval 2.5 μm, number of *z* slices varying depending on the thickness of organoids) were imaged using a NIDAQ *z* piezo and the widefield fluorescence imaging modality was used for brightfield detection with a Hamamatsu Orca Flash 4.0 camera at 100 ms exposure. For combined BF and POI-Halo organoid images (Fig. 1D, Fig. 2B), acquisition was performed using an Apo LWD 40× WI λS DIC N2 water immersion objective (N.A. 1.15; Nikon). To detect POI-Halo-JF635 fluorescence, JF635 was excited with 640 nm (laser at 40-80% depending on POI-Halo expression level) and detected with a Hamamatsu Orca Flash 4.0 camera at 200 ms exposure. Consecutively, the widefield fluorescence imaging modality was used for brightfield detection with a Hamamatsu Orca Flash 4.0 camera at 100 ms exposure. For each modality, multiple *z* planes (*xy* pixel size: 0.1625 × 0.1625 μm; *z* interval 0.4 μm, number of *z* slices varying depending on the thickness of organoids) were imaged using a NIDAQ *z* piezo. For TetON POI-Halo organoid images (Fig. S6A), only the POI-Halo-JF635 expression was recorded as described above.

#### Generation of 2D enteroid monolayer cultures from 3D mouse small intestinal organoids and seeding on imaging dishes

2D EMCs were derived from 3D mSIOs as described^15^ with the following specifications/exceptions: A 1:40 (vol:vol) mixture of Matrigel Growth Factor Reduced Basement Membrane Matrix (Corning, cat.# 356231) in DMEM/F-12 without phenol red was used for coating imaging dishes. Coated imaging dishes were incubated for at least 30 min up to one week in the cell culture incubator prior to seeding. For cell seeding, mSIO-derived single-cell suspensions with a concentration of 1×10^6^ cells/mL were prepared and 100-500 μL were seeded in plating medium (IntestiCult containing 20 μM ROCK inhibitor and 3 μM CHIR) to achieve a total of 500 μL/well of an 8-well Labtek II #1.5. Medium was exchanged to IntestiCult 1-2 d after seeding and exchanged every other day.

#### Confocal imaging of POI-Halo enteroid monolayer cultures

2D EMCs were seeded and cultured as described above. 5 d post-seeding, monolayers were stained with 200 nM HTL-JFX549 (kind gift from Luke Lavis) in DMEM/F-12 without phenol red for 30 min. Following two washes with DMEM/F-12 without phenol red for 15 min each, medium was replaced one more time to DMEM/F-12 without phenol red before proceeding with imaging.

Confocal imaging of live 2D EMCs (Fig. 1D, Fig. 2B) was performed on an LSM900 Airyscan 2 laser-scanning microscope with an inverted Axio Observer.Z1 / 7 operated by the ZEN 3.1 blue software (ZEISS). The microscope was equipped with a temperature- and CO_2_-controlled incubation chamber (Zeiss/PeCon), and the temperature was set to 37°C and the CO_2_ to 5%. Images were acquired using a Plan-Apochromat 40×/N.A. 1.3 Oil DIC (UV) VIS-IR M27 oil-immersion objective (ZEISS). One *z* plane was imaged in the 561 nm channel (*xy* pixel size: 0.092 × 0.092 μm; 1.21 μs pixel dwell time; bidirectional scanning; 4-times averaging). JFX549 was excited with 561 nm (diode (SH) laser at 0.1-3.0% depending on the POI-Halo expression level) and detected with GaAsP (spectral gallium arsenide) detectors at 566-635 nm.

#### IF of 2D enteroid monolayer cultures and confocal imaging

2D EMCs were seeded and cultured as described above. 5 d post-seeding, monolayers were fixed with 4% paraformaldehyde (PFA; Electron Microscopy Sciences, cat.# EMS14710) in PBS for 20 min at RT, washed three times with PBS for 5 min and stored at 4°C or directly subjected to IF staining. Cells in fixed and washed EMCs were permeabilized with 0.5% Triton X-100 (TX-100; Sigma-Aldrich, cat.# T9284) in PBS for 1 h. Following three washes for 5 min each with PBS, blocking was performed in 3% (vol/vol) donkey serum (Sigma-Aldrich, cat.# D9663) or 3% (vol/vol) goat serum (Sigma-Aldrich, cat.# G9023) in 0.1% (vol/vol) TX-100 in PBS (blocking buffer) for 4 h at RT or ON at 4°C. Incubation with the primary antibody (AB) in blocking buffer was performed ON at 4°C in a humidified chamber. Following three washes in blocking buffer for 5 min each, incubation with the secondary AB was performed for 1 h at RT. Following three washes with PBS for 5 min each, DNA was stained with 1 μg/mL Hoechst 33342 (Thermo Scientific, cat.# H3570) in PBS for 10-15 min, followed by another three washes with PBS for 5 min each. Immunostained samples were either imaged directly or sealed with parafilm and stored for short-term at 4°C prior to imaging.

The following primary ABs and dilutions were used: rabbit anti-Hes1 (Abcam, cat.# ab108937; 1:200), rabbit anti-Hnf4g (Sigma-Aldrich, cat.# HPA005438; 1:50), mouse anti-V5-tag (ThermoFisher, cat.# R960-25; 1:100), rabbit anti-Olfm4 (Cell Signaling, cat.# 39141; 1:50), rabbit anti-Aqp1 (Proteintech, cat.# 20333-1-AP; 1:200), rabbit anti-Ki67 (Abcam, cat.# ab16667; 1:200), sheep anti-Dll1 (Biotechne, cat.# AF3970; 1:20), rabbit anti-Lysozyme (Agilent, cat.# A009902-2; 1:300), rabbit anti-Aldolase B/C (Abcam, cat.# ab75751; 1:50).

Secondary donkey anti-rabbit-AlexaFluor568 (Thermo Scientific, cat.# A10042) or donkey anti-sheep-AlexaFluor568 ABs (Thermo Scientific, cat.# A21099) were used except for the V5 condition, for which a goat anti-mouse-AlexaFluor647 AB (Thermo Scientific, cat.# A21235) was used.

Confocal imaging of immunostained monolayers (Fig. S2C) was performed on an LSM900 Airyscan 2 laser-scanning microscope with an inverted Axio Observer.Z1 / 7 operated by the ZEN 3.1 blue software (ZEISS) at RT. Images were acquired using a Plan-Apochromat 40×/N.A. 1.3 Oil DIC (UV) VIS-IR M27 oil-immersion objective (ZEISS). One *z* plane was imaged in the 405 nm and 561 nm channels (*xy* pixel size: 0.11 × 0.11 μm; 2.06 μs pixel dwell time; bidirectional scanning; 4-times averaging). Hoechst was excited with 405 nm (diode laser at 0.2%) and detected with a multialkali-PMT (MA-PMT) detector at 410-493 nm. AlexaFluor568 was excited with 561 nm (diode (SH) laser at 0.5-5.0% depending on the brightness of the immunostaining) and detected with MA-PMT detector at 576-700 nm. AlexaFluor647 (for mouse anti-V5-tag only) was excited with 640 nm (diode laser at 3.0%) and detected with MA-PMT detector at 645-700 nm. Laser excitation powers were kept constant across all samples per immunostaining condition.

#### Image analysis of immunostained 2D enteroid monolayer cultures

Confocal images of immunostained 2D EMCs (Fig. 4C) were subjected to a Gaussian blur in the Hoechst channel, following which nuclei were segmented using the Cellpose^88^ Python package and the pre-trained default ‘cyto’ model, with the ‘diameter’ parameter set to 100 px (11.14 µm). The scikit-image library^89^ was then used to compute the morphological characteristics (nuclear area and eccentricity) of the masks generated by Cellpose. The nuclear masks were applied to the Ki67 channel and mean Ki67 intensities were extracted using the scikit-image library^89^.

#### Preparation of 2D enteroid monolayer cultures for SMT and PAPA-SMT

2D EMCs were seeded and cultured as described above. 2-6 d post-seeding (depending on the degree of monolayer confluency and the formation of crypt- and villus-like morphological characteristics), monolayers were stained for SMT with 50 nM HTL-JFX549 (bulk labeling for nuclear segmentation and feature extraction; kind gift from Luke Lavis) and 1 nM HTL-JFX650 (sparse labeling for SMT; kind gift from Luke Lavis) in DMEM/F-12 without phenol red for 15 min. Following two washes with DMEM/F-12 without phenol red for 15 min each, medium was replaced one more time to DMEM/F-12 without phenol red before proceeding with imaging. For SMT of TetON POI-Halo, 200 μg/mL doxycycline was added to the staining, washing, and imaging media. For PAPA-SMT, monolayers were stained with 50 nM HTL-JFX549 (sender) and 3 nM (Hes1) or 1 nM (Hnf4g) HTL-JFX650 (receiver) in DMEM/F-12 without phenol red for 15 min. Washes were performed as described for SMT.

#### TIRF microscope for HILO-based fast SMT and PAPA-SMT

All SMT and PAPA-SMT experiments were performed using a custom-built microscope as previously described^46^. In brief, a Nikon TI microscope was equipped with a 100×/N.A. 1.49 oil-immersion TIRF objective (Nikon apochromat CFI Apo TIRF 100× Oil), a motorized mirror, a perfect focus system, an EM-CCD camera (Andor iXon Ultra 897), a laser launch with 405 nm (140 mW, OBIS, Coherent), 488 nm, 561 nm and 639 nm (1 W, Genesis Coherent) laser lines, and an incubation chamber maintaining a humidified atmosphere with 5% CO_2_ at 37°C. All microscope, camera and hardware components were controlled through the NIS-Elements software (Nikon). Laser power densities used for imaging were approximately 52 W/cm^2^ for 405 nm (violet), 100 W/cm^2^ for 561 nm (green), and 2.3 kW/cm^2^ for 639 nm (red).

#### Microscope automation for SMT and PAPA-SMT

A custom-built inverted Nikon TI microscope described above was automated using code written in Python and the NIS Elements Macro Language. The microscope stage was moved to raster over a grid on the surface of the imaging dish. At each point on the grid, a 512 px x 512 px (6710.9 µm^2^) image was recorded in the densely labeled JFX549 channel. Nuclei were segmented using the pre-trained Versatile Fluorescent Nuclei model of the Python package StarDist^58^. One of the nuclei within a range of user-defined brightness and size parameters was chosen randomly, and the stage was moved to center it in the large FOV. The FOV was then resized to encompass a smaller 150 px x 150 px (576 µm^2^) square centered on the chosen nucleus. Images of the large and small FOV were recorded in multiple channels corresponding to the fluorophores in the sample (JFX549, JFX650). JFX650 fluorophores in the small FOV were pre-bleached with red light at 639 nm to achieve sparse and thus trackable single molecules, and an illumination sequence was executed for fast SMT or PAPA-SMT, respectively. Code for microscope automation will be made available upon acceptance of this manuscript for publication in a peer-reviewed journal.

#### Fast SMT of POI-Halo in enteroid monolayer cultures

For imaging in JFX549 or JFX650 channels, laser powers were set to 110 mW for 561 nm and to 1100 mW for 639 nm. The exposure time was set between 50 ms and 200 ms with a laser excitation power between 1.35% and 20% depending on the POI-Halo expression level to achieve segmentable bulk labeling and to avoid saturation. Conditions were kept constant across experiments for the same POI-Halo. Semrock 593/40 nm or 676/37 nm bandpass filters, respectively, were used.

For fast SMT experiments, the following bleaching durations in the 639 nm channel (100% laser excitation power) prior to executing an illumination sequence were used: NLS – 10s, H2B – 15s, Hes1 – 7s or 10s, Hnf4g – 10s, TetOn Hes1 – 10s, TetON Hnf4g – 20s.

The triggered illumination sequence consisted of the following phases at a frame rate of 7.48 ms/frame:

1. Imaging: 5000 frames of red light (639 nm, one 2 ms stroboscopic pulse per frame) with 5% 405 nm reactivation (pulsed during the camera transition time between 7 ms detection windows)
2. Imaging: 5000 frames of red light (639 nm, one 2 ms stroboscopic pulse per frame) with 10% 405 nm reactivation (pulsed during the camera transition time between 7 ms detection windows)

Red illumination was restricted to stroboscopic pulses during each frame of a single-molecule movie to reduce the motion blur of moving molecules^90^.

#### PAPA-SMT of POI-Halo in enteroid monolayer cultures

Imaging for PAPA-SMT was performed as described for fast SMT.

For PAPA-SMT experiments, the following bleaching durations in the 639 nm channel (100% laser excitation power) prior to executing an illumination sequence were used: Hes1 – 10s, Hnf4g – 10s.

For PAPA experiments, the illumination sequence consisted of 5 cycles of the following phases at a frame rate of 7.48 ms/frame:

1. Imaging: 250 frames of red light (639 nm), one 2 ms stroboscopic pulse per frame, recorded
2. DR pulse: 10 frames of violet light (405 nm), continuously during 7 ms detection window, recorded
3. DR imaging: 250 frames of red light, one 2 ms stroboscopic pulse per frame, recorded
4. Bleaching: 200 frames of red light, continuously during 7 ms detection window, not recorded
5. Imaging: 250 frames of red light, one 2 ms stroboscopic pulse per frame, recorded
6. PAPA pulse: 100 frames of green light (561 nm), continuously during 7 ms detection window, recorded
7. PAPA imaging: 250 frames of red light, one 2 ms stroboscopic pulse per frame, recorded
8. Bleaching: 200 frames of red light (639 nm), continuously during 7 ms detection window, not recorded

#### SMT and PAPA-SMT data processing and analysis

##### Image analysis for SMT and PAPA-SMT

Nuclei in epifluorescence images corresponding to SMT or PAPA-SMT movies were segmented using StarDist^58^ and then subjected to manual QC for downstream analysis using CellPicker, a MATLAB graphical user interface (GUI) that allows the user to filter nuclei based on their intensities in each channel and to perform QC to exclude nuclei with erroneous segmentation masks, unusual textures, or masks corresponding to segmented autofluorescent debris (https://github.com/tgwgraham/basic_PAPASMT_analysis). The morphological parameters nuclear area and eccentricity, as well as POI intensity in different channels as a proxy for relative POI expression level were computed for each of the chosen nuclei using the Python library scikit-image^89^ and the nuclear masks generated by StarDist.

##### SMT data processing and analysis

SMT movies were processed using quot (https://github.com/alecheckert/quot; ^59^), an open-source Python package that identifies single-molecule localizations and generates trajectories, and the following settings: [filter] start = 0; method = ‘identity’; chunk_size = 100; [detect] method = ‘llr’; k=1.0; w=9, t=18; [localize] method = ‘ls_int_gaussian’, window size = 9; sigma = 1.0; ridge = 0.001; max_iter = 10; damp = 0.3; camera_gain = 109.0; camera_bg = 470.0; [track] method = ‘conservativeeuclidean’; pixel_size_µm = 0.160; frame interval = 0.00748; search radius = 1; max_blinks = 0; min_IO = 0; scale – 7.0. A conservative method, in which only trajectories with unambiguously assigned localizations within a search radius of 1 µm in consecutive frames were considered, was used to construct trajectories. Trajectories were assigned to cells based on the StarDist-generated nuclear masks. Diffusion coefficient distributions were obtained using the state array method of the Python package saSPT (https://github.com/alecheckert/saspt; ^59^). Briefly, a regular Brownian motion with a normally distributed, mean-zero localization error (RBME) model was fit to populations of single-molecule trajectories in each cell to obtain a state array, consisting of posterior occupations of states defined by their localization errors and diffusion coefficients. These state array distributions were then marginalized on the diffusion coefficients to obtain diffusion spectra. Mean diffusion spectra for each protein were constructed by averaging over single-cell diffusion spectra, with cells weighted by the number of trajectories. The fraction of bound trajectories for a POI was defined as the fraction of trajectories with diffusion coefficients below 0.15 µm^2^/s in these mean diffusion spectra (fraction bound). The diffusion coefficient corresponding to the highest posterior occupation was defined as the modal diffusion coefficient. Fractions of bound trajectories and modal diffusion coefficients were also computed for individual nuclei based on the single-cell diffusion spectra to correlate them with other extracted cellular features, such as nuclear area and eccentricity, as well as POI expression level. A custom-written Jupyter notebook will be made available upon acceptance of this manuscript for publication in a peer-reviewed journal.

##### Cluster analysis based on single-cell diffusion spectra

For cluster-based analyses, a matrix of pairwise distances between pairs of single-cell diffusion spectra was computed using the Jensen-Shannon distance as the distance metric^62^. The AgglomerativeClustering class of the scikit-learn Python library^91^ was then used to hierarchically cluster cells with complete linkage. The over- and under-representation of cells expressing different POIs in different clusters was quantified in terms of a *p*-value computed from the hypergeometric distribution. The significance threshold was Bonferroni-corrected to account for multiple comparisons. A custom-written Jupyter notebook will be made available upon acceptance of this manuscript for publication in a peer-reviewed journal.

##### PAPA data processing and analysis

PAPA movies were processed and analyzed as described for SMT with the following specifications: Custom MATLAB code reported previously (^55^; https://github.com/tgwgraham/basic_PAPASMT_analysis) was used to extract all trajectory segments occurring within the first 30 frames after pulses of 561 nm light (PAPA trajectories) or 405 nm light (DR trajectories). PAPA and DR trajectories were then separately analyzed similar to SMT data.

### Quantification and statistical analysis

For SMT experiments, bootstrapping on all combined experiments per condition was performed by drawing 1000 samples from the population with replacement, with each sample containing as many nuclei as the total number of nuclei in the population under analysis. Diffusion spectra marginalized on the diffusion coefficients were generated for each bootstrap replicate, and these were used to compute confidence intervals for the fraction of bound trajectories. A custom-written Jupyter notebook will be made available upon acceptance of this manuscript for publication in a peer-reviewed journal.

For PAPA-SMT experiments, statistical analysis was performed as described before using custom-written MATLAB scripts (^47^; https://github.com/tgwgraham/basic_PAPASMT_analysis). For a side-by-side comparison of the distributions for DR and PAPA trajectories, they were randomly subsampled without replacement for the condition with more trajectories. Following subsampling, bootstrapping analysis with replacement was performed on all combined experiments per condition for the PAPA datasets for Hes1 and Hnf4g. For each combined dataset, a random sample of size *n*, where *n* is the total number of cells in the combined dataset, was drawn 100 times. The mean and standard deviation from these analyses is reported. For significance testing between DR and PAPA, two-tailed *p*-values were calculated based on a normal distribution (Scipy function, scipy.stats.norm.sf) with mean equal to the difference between sample means and variance equal to the sum of the variances from the bootstrap resampling.

### Additional resources

#### Data and code availability

Custom-written code is partially available via the indicated links to GitLab repositories and will be made fully available upon acceptance of this manuscript for publication in a peer-reviewed journal. Data reported in this manuscript will be made publicly available upon acceptance of this manuscript for publication in a peer-reviewed journal.

## Acknowledgements

We would like to thank current and past members of the Tjian/Darzacq lab for scientific discussions. We are grateful to Qiulin Zhu and Brendan Wu for cloning assistance, and Shuang Zheng for assistance with mouse colony maintenance. We thank Claudia Cattoglio for advice regarding lentivirus production. We are thankful to Thomas Graham for sharing code for SMT and PAPA automation and analysis as well as for discussions about PAPA data analysis and interpretation. We also thank Luke Lavis for providing JF dyes and the UC Berkeley Cell Culture Facility supported by The University of California, Berkeley, for providing cell lines. We are grateful to Ophir Klein and current and past members of the Klein lab gut group for sharing protocols, providing a platform for discussion and feedback throughout this study, and for critically commenting on the figures. We would like to thank Franziska Lorbeer for critically reading and commenting on the manuscript. N.W. acknowledges funding from the Berkeley Stem Cell Center via a Siebel postdoctoral fellowship and from the German Research Foundation (DFG) via a Walter Benjamin fellowship (453309976). S.A. was supported by a Donner 160 fellowship. R.T. acknowledges funding from the Howard Hughes Medical Institute (34430) and X.D. from the Chan Zuckerberg Initiative via a Dynamic Imaging Grant (Dynamic-0000000091).

## Author contributions

N.W. conceptualized and designed the study, executed all experiments, supervised the development of the automated imaging and analysis pipeline, analyzed data, and wrote the manuscript (original draft, review/editing). S.A. developed the automation pipeline and tailored SMT analysis code including cellular feature extraction, cluster, and statistical analyses, segmented confocal data, discussed data and their interpretation with N.W., provided a subset of raw figure panels as well as methods details, and reviewed the manuscript. G.M.D. designed and cloned DNA constructs. R.T. and X.D. hosted the study, provided equipment, and reviewed the manuscript. N.W., R.T., and X.D. provided funding for this project. All authors agreed on the final version of the manuscript.

## Inclusion and diversity

N.W., S.A., and G.M.D. support inclusive, diverse, and equitable conduct of research.

