## Supplemental figure legends for "Automated live-cell single-molecule tracking in enteroid monolayers reveals transcription factor dynamics probing lineage-determining function"

**Supplemental information**

Supplemental figures S1-S12

### Supplemental figure legends

<u>Figure S1:</u> **Fast SMT of H2B-Halo and Halo-NLS as slowly and fast diffusing controls in 2D EMCs, related to Figure 1. (A)** Single-cell diffusion heatmaps for 3 combined experiments each for H2B (left; red) and NLS (right; gray). Cells are ordered according to increasing fraction bound from bottom to top. **(B)** Single-cell diffusion spectra for 3 independent experiments each for H2B (left; red) and NLS (right; gray). Mean diffusion spectra are plotted in black (bold line), and the fraction bound is indicated. **(C)** Single-cell cumulative distribution functions (CDFs) for 3 independent experiments each for H2B (left; red) and NLS (right; gray). The mean CDFs for the 3 combined experiments are plotted in black (bold line). Cells in (A-C) correspond to the 3 experiments each for H2B (red; combined *n*=127 cells) and NLS (gray; combined *n*=92 cells) with mean diffusion spectra plotted in Fig. 1G.

<u>Figure S2:</u> **Characterization of 3D enteroid lines and derived 2D EMCs stably expressing Hes1-Halo and Hnf4g-Halo, related to Figure 2. (A,B)** BF overview images of (A) Hes1-Halo or (B) Hnf4g-Halo organoids displaying a budding morphology 5 d post-seeding. Scale bars: 200 µm. **(C)** Immunofluorescence experiments probing for POI and cell type markers (magenta) in 2D EMCs co-stained with Hoechst (gray) 5 d after their derivation from wild-type (WT; left), Hes1-Halo (middle), or Hnf4g-Halo (right) 3D enteroids. White asterisks indicate Dll1-positive cells. Experiments in (A,B) were performed two months after the establishment of stable mSIO lines. Scale bars: 20 µm.

<u>Figure S3:</u> **Single-cell diffusion spectra of a manual fast SMT experiment of Hes1-Halo in 2D EMCs, related to Figure 2. (A)** Single-cell diffusion spectra for Hes1 (light turquoise). The mean diffusion spectrum is plotted in black (bold line), and the fraction bound is indicated. **(B)** Single-cell CDFs for Hes1 (light turquoise). The mean CDF is plotted in black (bold line). Cells correspond to the manual experiment (*n*=15 cells) with the single-cell diffusion heat map plotted in Fig. 2C.

<u>Figure S4:</u> **Automated fast SMT of H2B-Halo in 2D EMCs, related to Figure 2. (A)** Single-cell diffusion heatmap for 5 combined automated experiments for H2B-Halo (combined *n*=1022 cells with 134,90,92,355,351 cells per experiment). Cells are ordered according to increasing fraction bound from bottom to top. **(B)** Numbers for H2B-Halo cells imaged, positively QCed, and subjected to correlative diffusion analysis and cellular feature extraction in the combined automated experiments from (A). **(C)** Single-cell diffusion spectra for H2B (red) with mean diffusion spectrum plotted in black (bold line) and fraction bound indicated. Bootstrap analysis of 5 combined experiments determined a fraction bound of 78.7% (95% CI: 76.8-80.6%). **(D)** Single-cell CDFs for H2B (red) with mean CDF plotted in black (bold line).

<u>Figure S5:</u> **Automated fast SMT of Hes1-Halo and Hnf4g-Halo in 2D EMCs, related to Figure 2. (A)** Single-cell diffusion spectra for Hes1 (light turquoise; left) and Hnf4g (dark turquoise; right) with mean diffusion spectra plotted in black (bold line) and fractions bound indicated. Bootstrap analysis of 8 combined automated experiments with *n*=9,15,17,32,64,33,60,37,68 cells for Hes1 determined a mean fraction bound of 31.7% (95% CI: 26.3-37.3%) and 3 combined automated experiments with *n*=68,18,100 cells for Hnf4g determined a mean fraction bound of 56.0% (95% CI: 52.4-58.9%). **(B)** Single-cell CDFs for Hes1 (light turquoise; left) and Hnf4g (light turquoise; right) with mean CDFs

plotted in black (bold line). **(C)** Violin plot for the fraction bound distributions (median (white point); first/third quartile (whiskers)) for Hes1 (light turquoise) and Hnf4g (dark turquoise) of the combined automated experiments from (A,B). Cells correspond to the experiments with single-cell diffusion heatmaps plotted in Fig. 2E and combined mean diffusion spectra plotted in Fig. 2F.

**Figure S6:** **Fast SMT of Hes1-Halo and Hnf4g-Halo upon induced expression in 2D EMCs, related Figure 2. (A)** Representative confocal images of Hes1-Halo (left) and Hnf4g-Halo (right) 1 d after doxycycline-induced expression (TF (gray)) in 3D enteroids 5 d post-seeding. Scale bars: 50 µm. **(B)** Single-cell diffusion heatmaps for 4 combined automated fast SMT experiments for TetOn Hes1 (left; light turquoise; combined *n*=141 cells with 52,17,15,57 cells per experiment) and for 4 combined automated fast SMT experiments for TetON Hnf4g (right; dark turquoise; combined *n*=213 cells with 26,11,63,113 cells per experiment) 1 d after doxycycline-induced transgene expression. Cells are ordered according to increasing fraction bound from bottom to top. **(C)** Single-cell diffusion spectra for TetOn Hes1 (light turquoise; left) and TetOn Hnf4g (dark turquoise; right) with mean diffusion spectra plotted in black (bold line) and fractions bound indicated. Bootstrap analysis of 4 combined experiments with *n*=141 cells for Hes1 determined a mean fraction bound of 33.9% (95% CI: 29.7-38.7%) and 4 combined experiments with *n*=213 cells for Hnf4g determined a mean fraction bound of 46.9% (95% CI: 42.4-51.4%). **(D)** Single-cell CDFs for TetON Hes1 (light turquoise; left) and TetON Hnf4g (light turquoise; right) with mean CDFs plotted in black (bold line).

**Figure S7:** **Comparative diffusion behavior of NLS, Hes1, Hnf4g, and H2B in 2D EMCs, related to Figure 3. (A)** Mean diffusion spectra and **(B)** mean CDFs for NLS (gray), Hes1 (light turquoise), Hnf4g (dark turquoise), and H2B (red) corresponding to the single-cell data shown in Fig. 3A. **(C)** Joint hierarchical clustering of Hes1, Hnf4g, H2B, and NLS based on single-cell diffusion spectra using the Jensen-Shannon distance metric referring to Fig. 3B. Top: Cluster statistics. Bottom left: *p*-values indicating the representation of each POI (NLS (cross), Hes1 (circle), Hnf4g (square), H2B (ex mark)) in each diffusion cluster (1 (blue), 2 (orange), 3 (green)) relative to the representation in the population; Bonferroni-corrected significance threshold (dashed line). Bottom right: Heatmap of *p*-values indicating the representation of each POI in each diffusion cluster (red: overrepresentation; blue: underrepresentation). **(D)** Joint hierarchical clustering of Hes1 and Hnf4g based on single-cell diffusion spectra using the Jensen-Shannon distance metric referring to Fig. 3C. Top: Cluster statistics. Bottom left: *p*-values indicating the representation of each POI (Hes1 (circle), Hnf4g (ex mark)) in each diffusion cluster (1 (blue), 2 (orange), 3 (green), 4 (red), 5 (purple)) relative to the representation in the population; Bonferroni-corrected significance threshold (dashed line). Bottom right: Heatmap of *p*-values indicating the representation of each POI in each diffusion cluster (red: overrepresentation; blue: underrepresentation).

**Figure S8:** **Cell level-based correlation of fast SMT-derived diffusion parameters of Hes1 and Hnf4g with cellular morphological characteristics in 2D EMCs, related to Figure 4. (A)** Top: Single-cell diffusion spectra for Hes1 color-coded according to diffusion clusters obtained through hierarchical clustering using the Jensen-Shannon distance metric. Bottom: Cluster statistics. **(B)** Single-cell pairplots of diffusion parameters (fraction bound, modal diffusivity) and cellular morphological characteristics (nuclear size, eccentricity) for Hes1 including histograms for each parameter (diagonal from top left to bottom right) with color-coded diffusion clusters (1 (blue), 2 (orange), 3 (green), 4 (red), 5 (purple), 6 (brown)). **(C)** Top: Single-cell diffusion spectra for Hnf4g color-coded according to diffusion clusters obtained through hierarchical clustering using the Jensen-Shannon distance metric. Bottom: Cluster statistics. **(D)** Single-cell pairplots of diffusion parameters and cellular morphological characteristics for Hnf4g including histograms for each parameter (diagonal from top left to bottom right) with color-coded diffusion clusters (1 (blue), 2 (orange), 3 (green)). Clustered Hes1 data in (A,B) are the same as in Fig. 4D. Clustered Hnf4g data in (C,D) are the same as in Fig. 4E.

**<u>Figure S9:</u> Cell level-based correlation of fast SMT-derived diffusion parameters of H2B with cellular morphological characteristics in 2D EMCs, related to Figure 4. (A)** Top: Single-cell diffusion spectra for H2B color-coded according to diffusion clusters obtained through hierarchical clustering using the Jensen-Shannon distance metric. Bottom: Cluster statistics. **(B)** Single-cell pairplots of diffusion parameters (fraction bound, modal diffusivity) and cellular morphological characteristics (nuclear size, eccentricity) for H2B including histograms for each parameter (diagonal from top left to bottom right) with color-coded diffusion clusters (1 (blue), 2 (orange), 3 (green)). Data are from 5 combined automated experiments for H2B-Halo (combined *n*=1022 cells with *n*=134,90,92,355,351 cells per experiment) and the same as plotted in Fig. S4.

**<u>Figure S10:</u> Cell level-based correlation of fast SMT-derived diffusion parameters for NLS, Hes1, Hnf4g, and H2B with cellular morphological characteristics in 2D EMCs, related to Figure 4.** Single-cell pairplots of diffusion parameters (fraction bound, modal diffusivity) and cellular morphological characteristics (nuclear size, eccentricity) for NLS (gray), Hes1 (light turquoise), Hnf4g (dark turquoise), and H2B (red) including histograms for each parameter (diagonal from top left to bottom right). Data correspond to the one plotted in Fig. 3A and Fig. S7.

**<u>Figure S11:</u> Detectability of molecular events by DR- *versus* PAPA-based SMT, related to Figure 5.** Molecular association events of a mixture of sender- (green) or receiver-labeled (red) Halo-POIs (gray-orange) detectable (yellow asterisk) or non-detectable (black asterisk) in fast SMT using DR (violet; top) *versus* PAPA (green; bottom) illumination through proximity of sender and receiver (gray cloud). i) Single monomeric POIs; ii) Dimerization (top) or lower-order oligomerization (bottom) of sender-/receiver-labeled molecules; iii) Larger-order oligomerization of sender-/receiver-labeled molecules; iv) Single monomeric POIs (randomly) in proximity; v) Single monomeric POIs spaced sufficiently apart.

**<u>Figure S12:</u> PAPA-SMT for studying self-association of Hes1 and Hnf4g in 2D EMCs, related to Figure 5. (A)** Mean number of localizations (black) detected in PAPA-SMT of double-labeled Hes1-Halo (left) or Hnf4g-Halo (right) consisting of five cycles of alternating DR (violet) and PAPA pulses (green). The mean localization number is averaged over all cells from 3 experiments each for Hes1 (232 cells total with *n*=84,90,58 cells per experiment) with 3058 subsampled trajectories or for Hnf4g (326 cells total with *n*=60,131,135 cells per experiment) with 12282 subsampled trajectories. **(B)** Mean number of localizations averaged over all five cycles of alternating DR (violet) and PAPA pulses (green) for fast PAPA-SMT of Hes1 (left) or Hnf4g (right) with 3058 or 12282 subsampled trajectories for Hes1 or Hnf4g, respectively. The mean localization number is plotted both as average for each of the three experiments (gray) and as total average across the three experiments (black). The first 30 frames following each light pulse used for analysis are colored in purple (DR) or green (PAPA). **(C)** Diffusion spectra for PAPA (green) *versus* DR (violet) resulting from bootstrap resampling with 100 replicates (one replicate per line) of 3 combined experiments for Hes1 (left) or Hnf4g (right). Data correspond to experiments plotted in Fig. 5C,D.
