## Supplemental figures for "Automated live-cell single-molecule tracking in enteroid monolayers reveals transcription factor dynamics probing lineage-determining function"

Figure S1

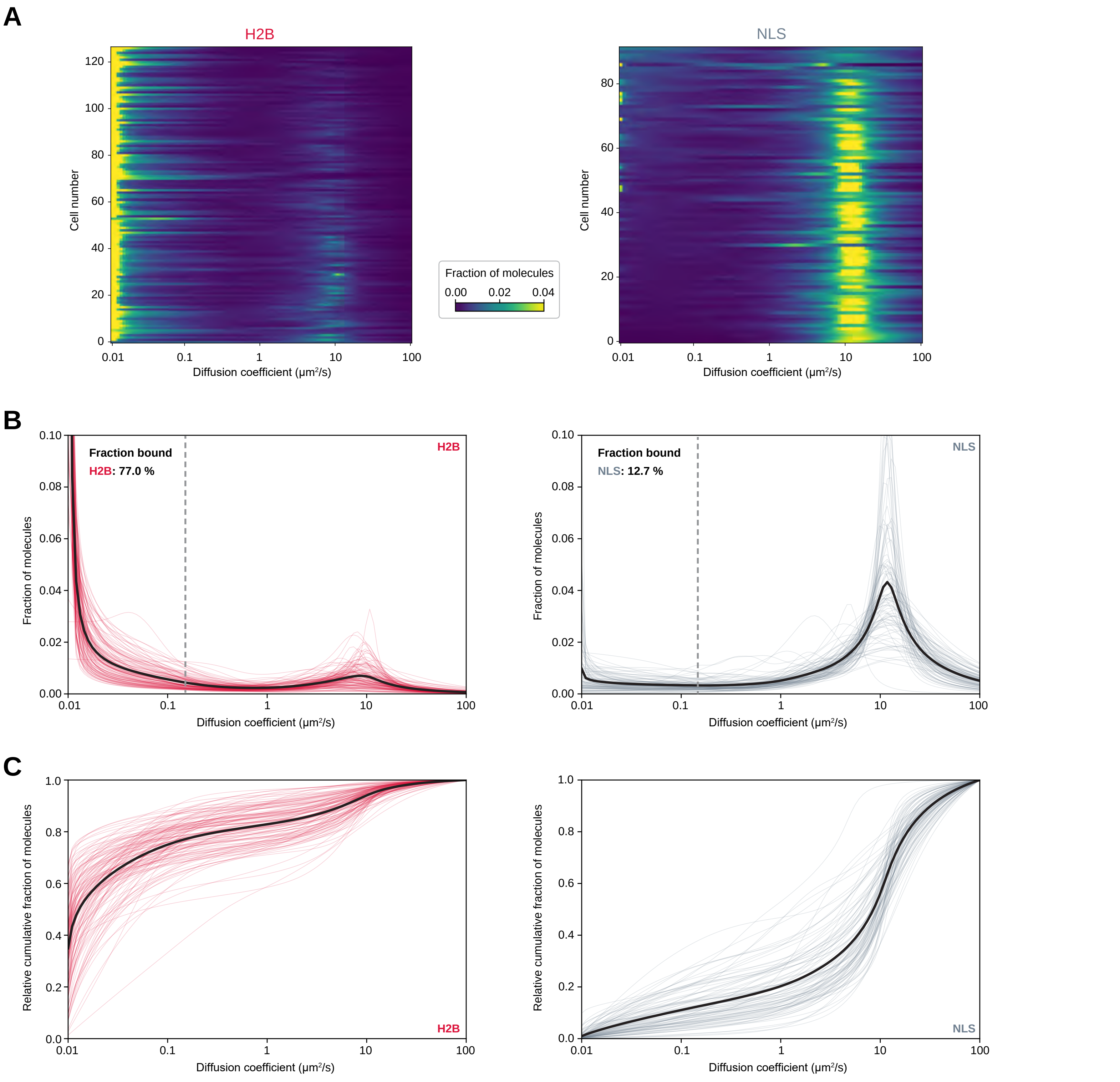
A
H2B
NLS
Cell number
Diffusion coefficient (μm²/s)
Fraction of molecules
0.00 0.02 0.04
B
Fraction bound
H2B: 77.0 %
NLS: 12.7 %
Fraction of molecules
Diffusion coefficient (μm²/s)
C
Relative cumulative fraction of molecules
Diffusion coefficient (μm²/s)
H2B
NLS

Figure S2

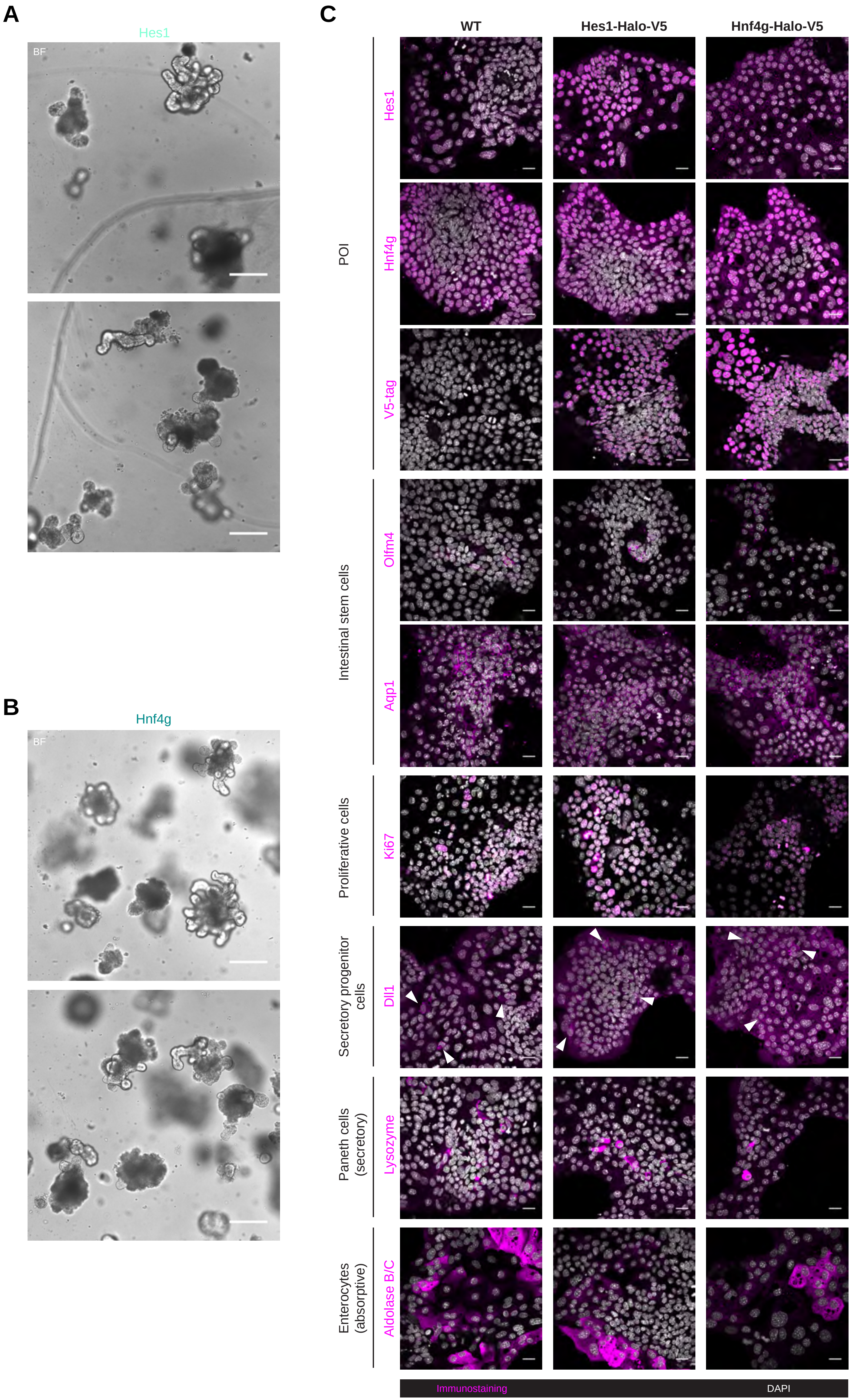

A
Hes1
BF
B
Hnf4g
BF
C
WT
Hes1-Halo-V5
Hnf4g-Halo-V5
POI
Hes1
Hnf4g
V5-tag
Intestinal stem cells
Olfm4
Aqp1
Proliferative cells
Ki67
Secretory progenitor cells
Dll1
Paneth cells (secretory)
Lysozyme
Enterocytes (absorptive)
Aldolase B/C
Immunostaining
DAPI

Figure S3

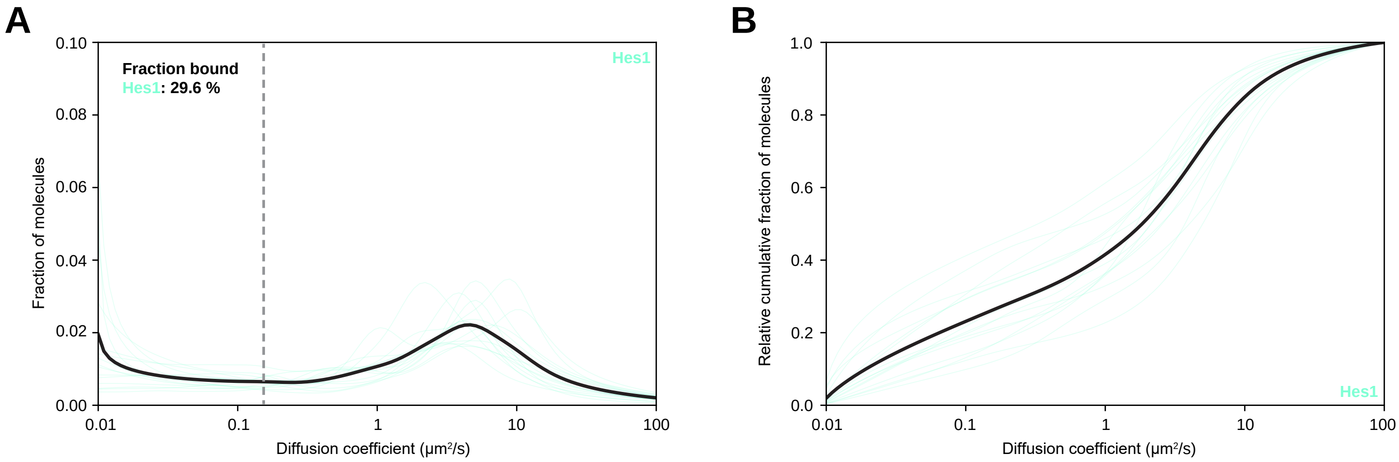
A
Fraction bound
Hes1: 29.6 %
Hes1
Fraction of molecules
0.10
0.08
0.06
0.04
0.02
0.00
0.01
0.1
1
10
100
Diffusion coefficient (μm²/s)
B
Relative cumulative fraction of molecules
1.0
0.8
0.6
0.4
0.2
0.0
Hes1
Diffusion coefficient (μm²/s)

Figure S4

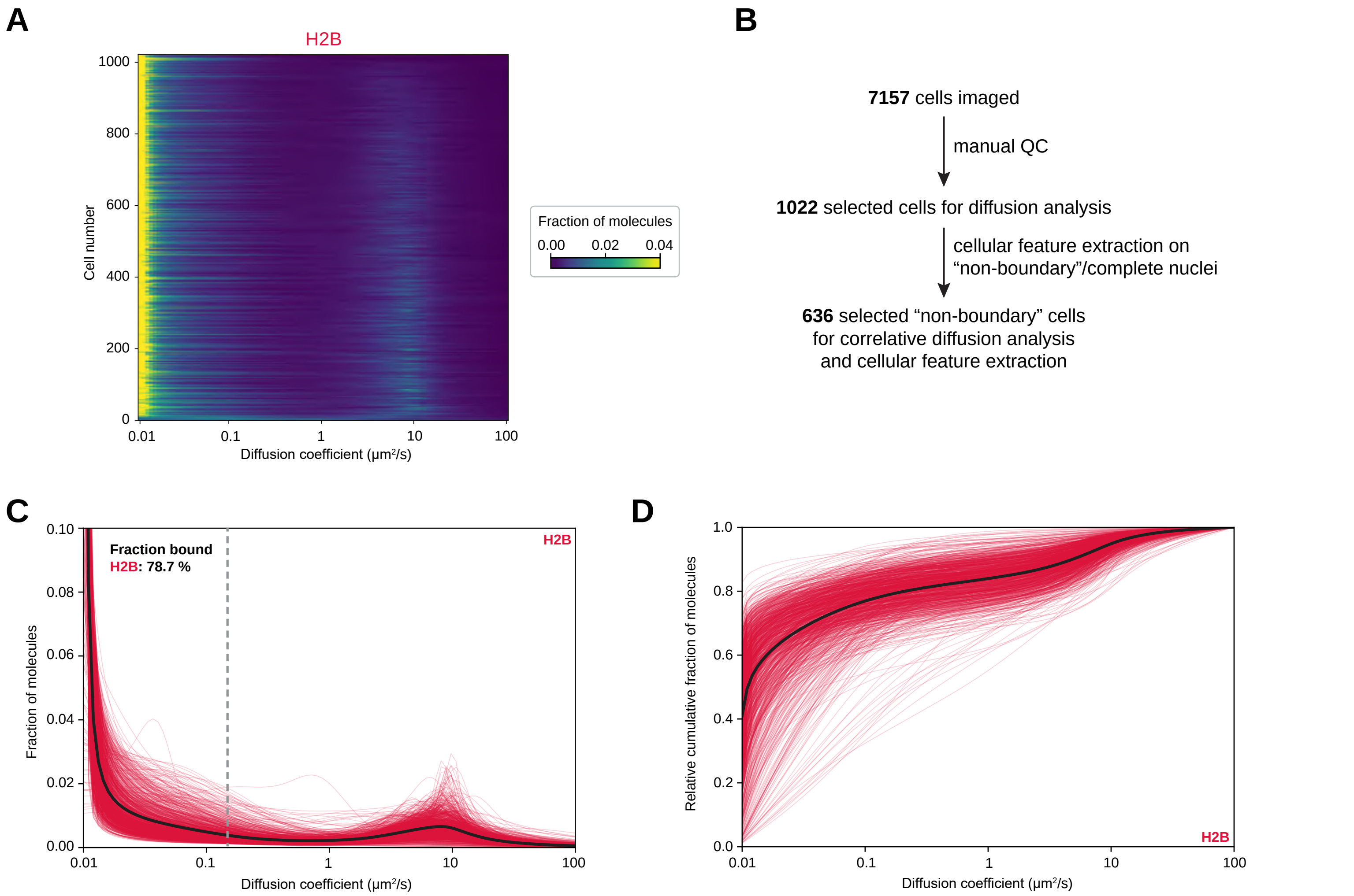
A
H2B
Cell number
1000
800
600
400
200
0
0.01
0.1
1
10
100
Diffusion coefficient (μm²/s)
Fraction of molecules
0.00
0.02
0.04
B
7157 cells imaged
manual QC
1022 selected cells for diffusion analysis
cellular feature extraction on
“non-boundary”/complete nuclei
636 selected “non-boundary” cells
for correlative diffusion analysis
and cellular feature extraction
C
Fraction bound
H2B: 78.7 %
H2B
Fraction of molecules
0.10
0.08
0.06
0.04
0.02
0.00
0.01
0.1
1
10
100
Diffusion coefficient (μm²/s)
D
Relative cumulative fraction of molecules
1.0
0.8
0.6
0.4
0.2
0.0
0.01
0.1
1
10
100
Diffusion coefficient (μm²/s)
H2B

Figure S5

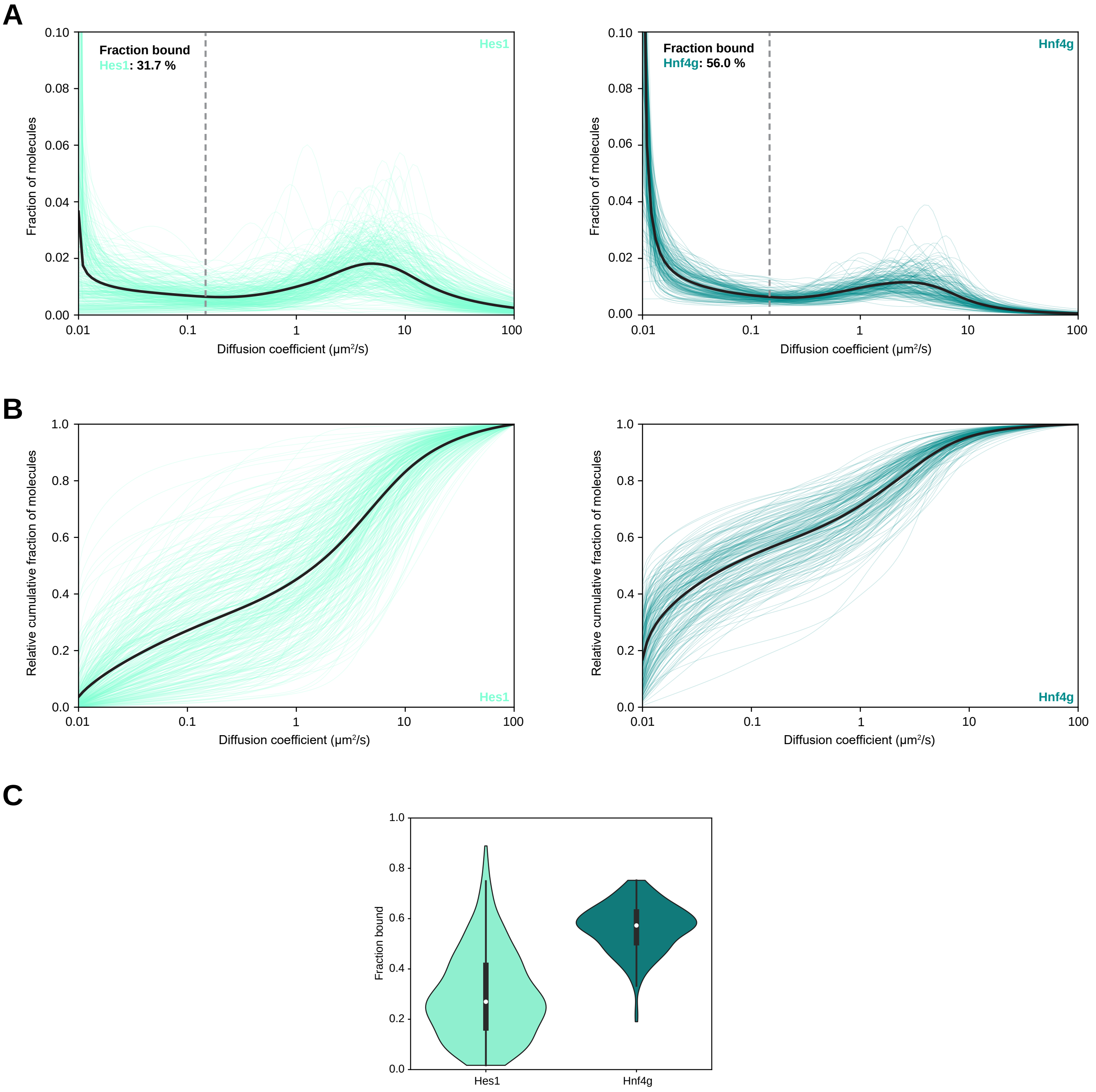
A
Fraction bound
Hes1: 31.7 %
Hes1
Fraction of molecules
0.10
0.08
0.06
0.04
0.02
0.00
0.01
0.1
1
10
100
Diffusion coefficient (μm²/s)
Fraction bound
Hnf4g: 56.0 %
Hnf4g
B
Relative cumulative fraction of molecules
1.0
0.8
0.6
0.4
0.2
0.0
Hes1
Hnf4g
Diffusion coefficient (μm²/s)
C
Fraction bound
Hes1
Hnf4g

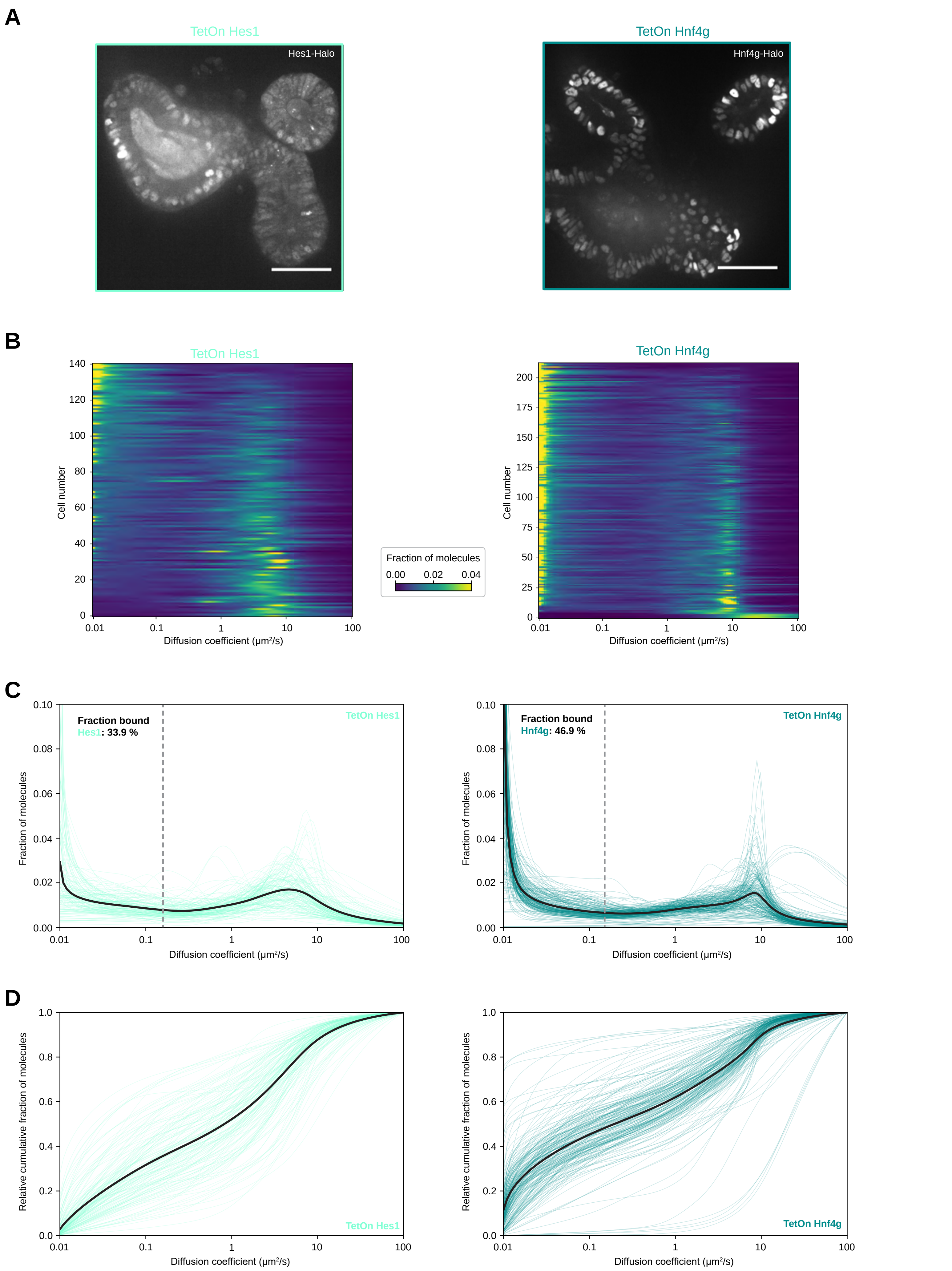
A
TetOn Hes1
Hes1-Halo
TetOn Hnf4g
Hnf4g-Halo
B
TetOn Hes1
Cell number
Diffusion coefficient (µm²/s)
Fraction of molecules
0.00 0.02 0.04
TetOn Hnf4g
Cell number
Diffusion coefficient (µm²/s)
C
Fraction bound
Hes1: 33.9 %
TetOn Hes1
Fraction of molecules
Diffusion coefficient (µm²/s)
Fraction bound
Hnf4g: 46.9 %
TetOn Hnf4g
Fraction of molecules
Diffusion coefficient (µm²/s)
D
Relative cumulative fraction of molecules
TetOn Hes1
Diffusion coefficient (µm²/s)
Relative cumulative fraction of molecules
TetOn Hnf4g
Diffusion coefficient (µm²/s)

Figure S7

**A**

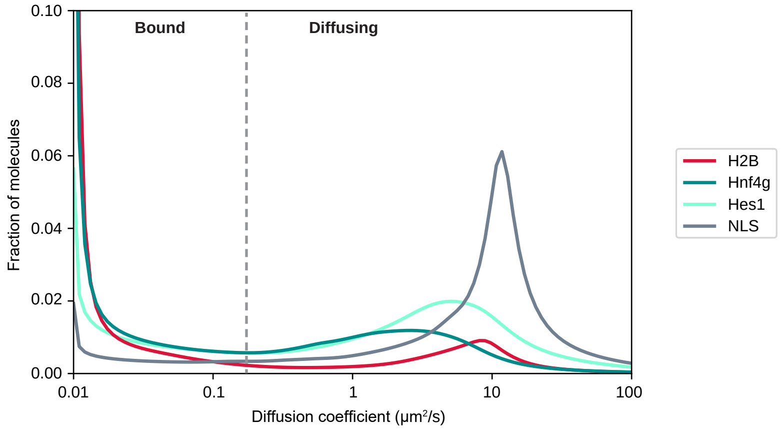


**B**

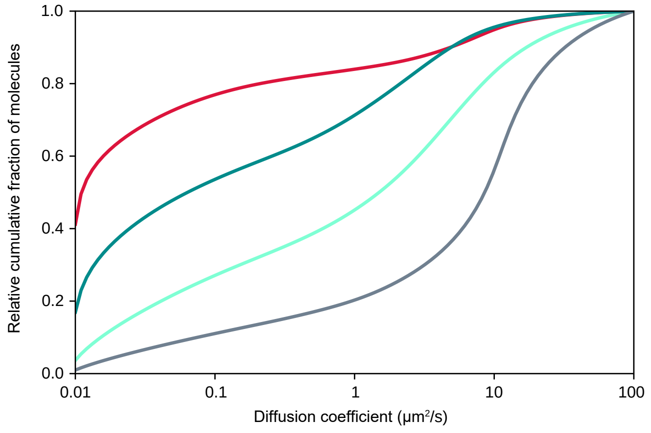


**C**

Joint clustering:
Hes1, Hnf4g, H2B, NLS

| Cluster | 1 | 2 | 3 |
|---|---|---|---|
| *f* bound | 76.7% | 37.9% | 12.0% |
| Modal *D* (μm²/s) | 0.01 | 0.01 | 11.77 |

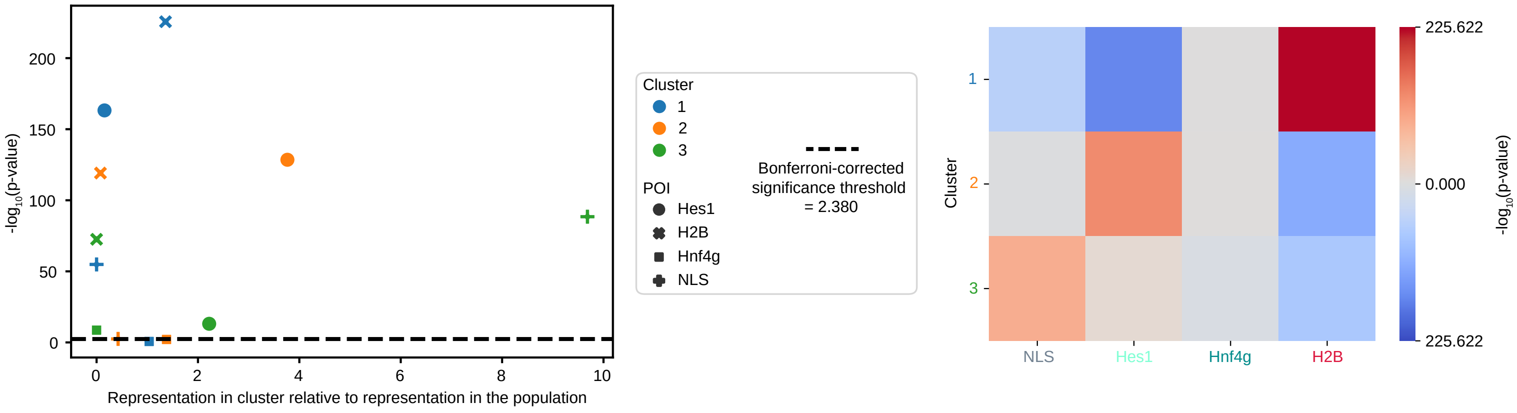


**D**

Joint clustering:
Hes1, Hnf4g

| Cluster | 1 | 2 | 3 | 4 | 5 |
|---|---|---|---|---|---|
| *f* bound | 50.0% | 34.1% | 57.3% | 12.6% | 15.0% |
| Modal *D* (μm²/s) | 0.01 | 0.01 | 0.01 | 8.11 | 4.23 |

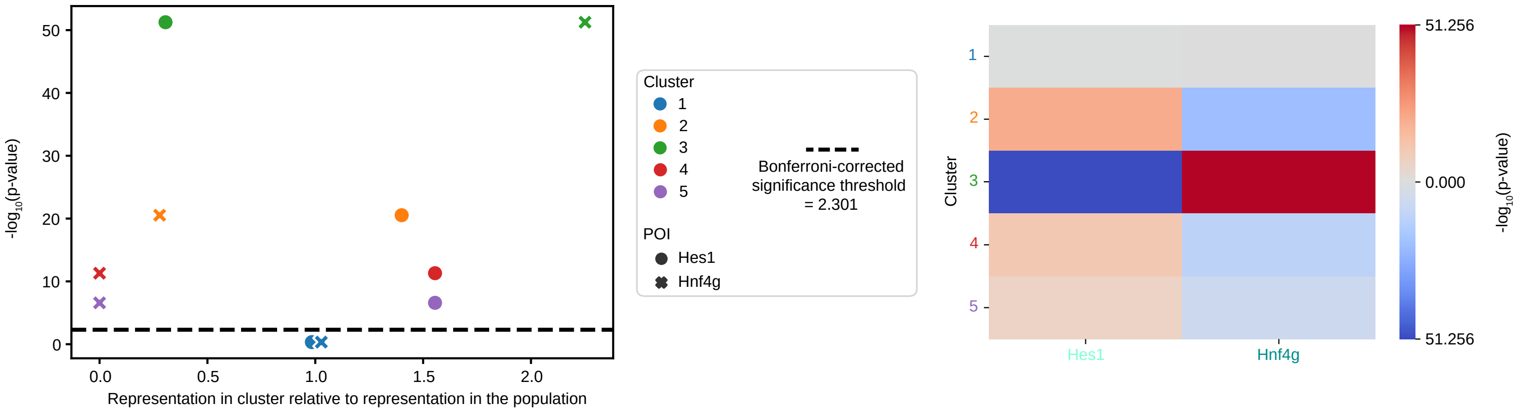

A

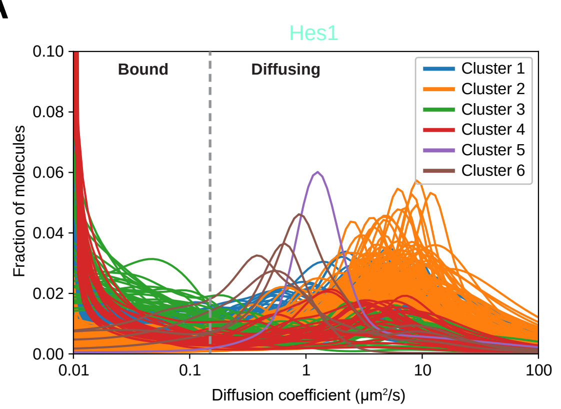


| Cluster | 1 | 2 | 3 | 4 | 5 | 6 |
|---|---|---|---|---|---|---|
| # Cells | 163 | 107 | 33 | 27 | 1 | 4 |
| % Cells | 48.7% | 31.9% | 9.9% | 8.1% | 0.3% | 1.2% |
| *f* bound | 32.0% | 14.5% | 56.3% | 53.6% | 2.7% | 21.4% |
| Modal *D* (µm²/s) | 0.01 | 7.39 | 0.01 | 0.01 | 1.26 | 0.79 |

B

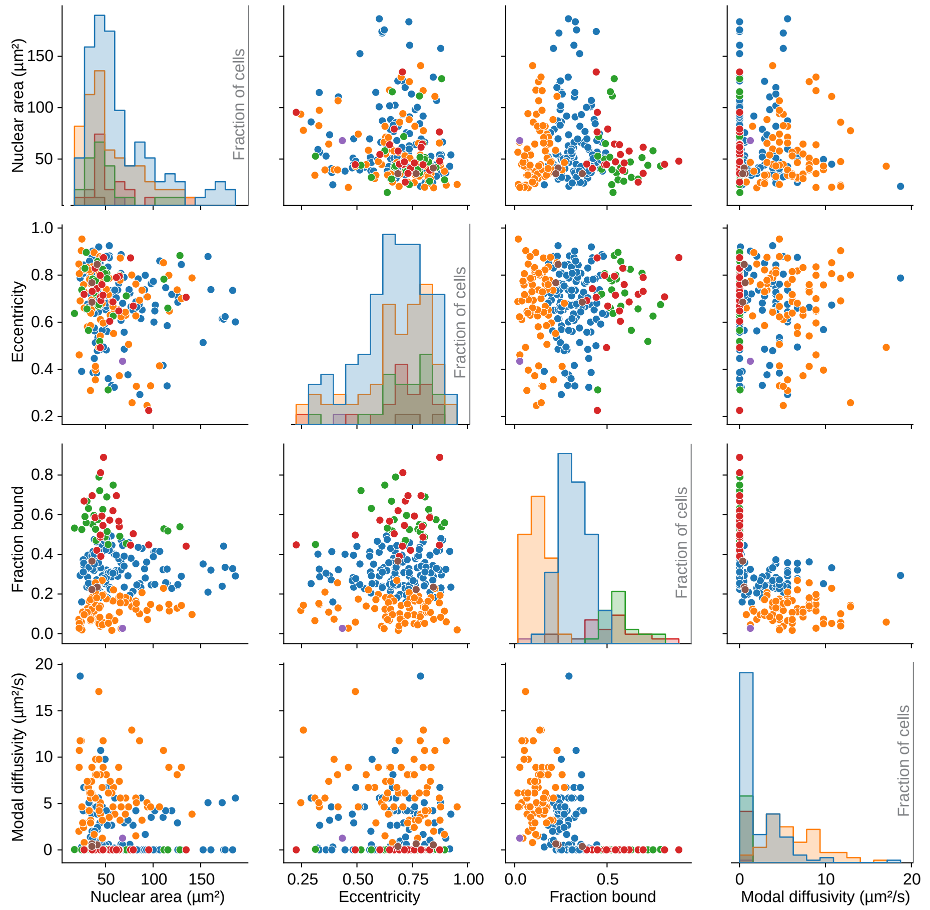


C

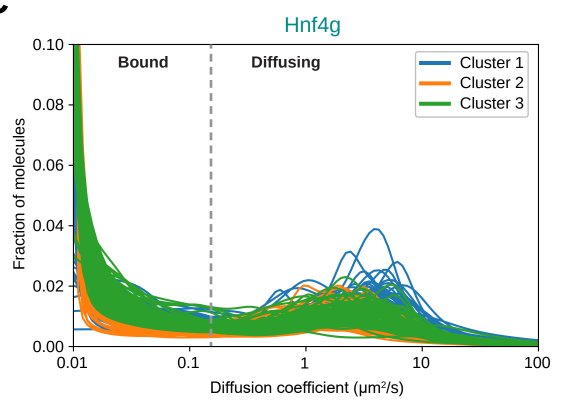


| Cluster | 1 | 2 | 3 |
|---|---|---|---|
| # Cells | 44 | 69 | 73 |
| % Cells | 23.7% | 37.1% | 39.2% |
| *f* bound | 43.4% | 60.4% | 56.2% |
| Modal *D* (µm²/s) | 0.01 | 0.01 | 0.01 |

D

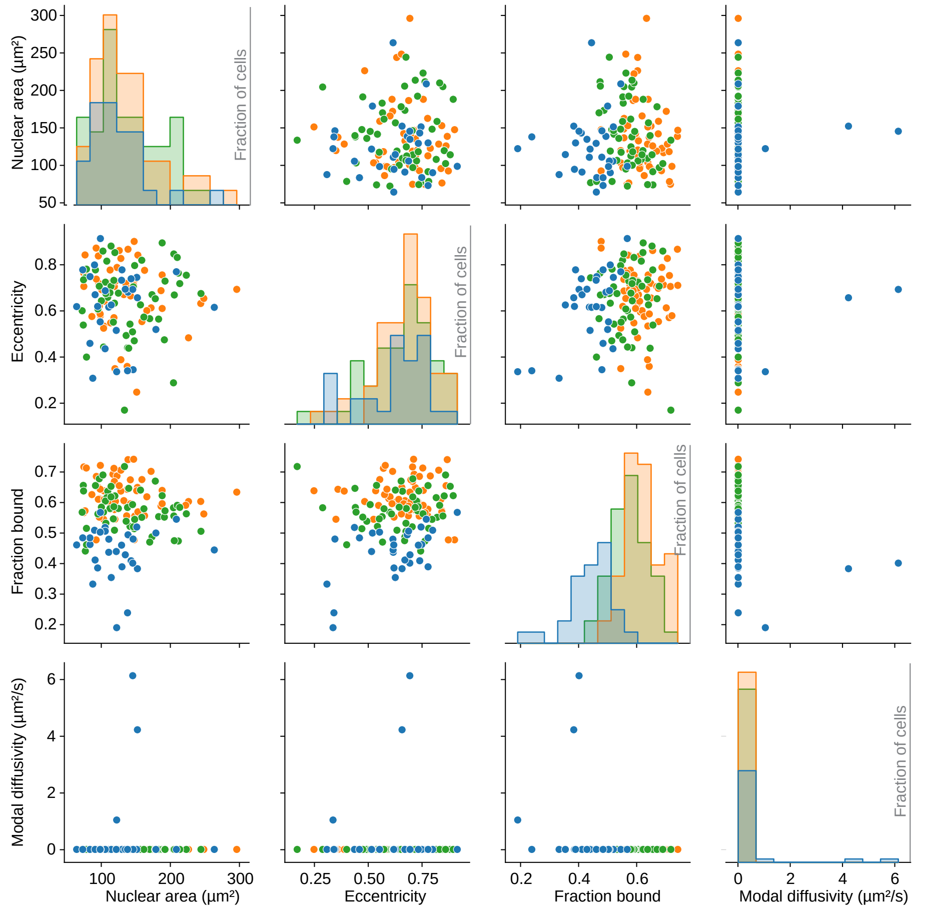

### Figure S9

**A**

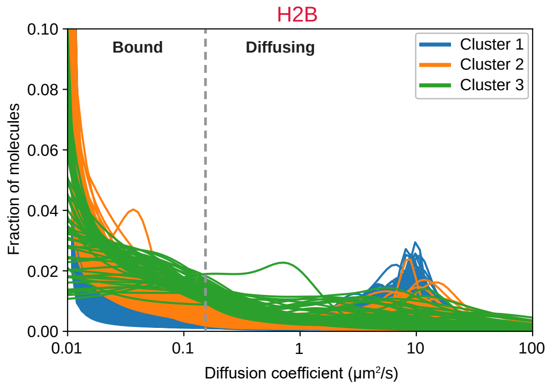


| Cluster | 1 | 2 | 3 |
|---|---|---|---|
| # Cells | 700 | 271 | 51 |
| % Cells | 68.5% | 26.5% | 5.0% |
| *f* bound | 78.7% | 78.8% | 68.9% |
| Modal *D* (µm²/s) | 0.01 | 0.01 | 0.01 |

**B**

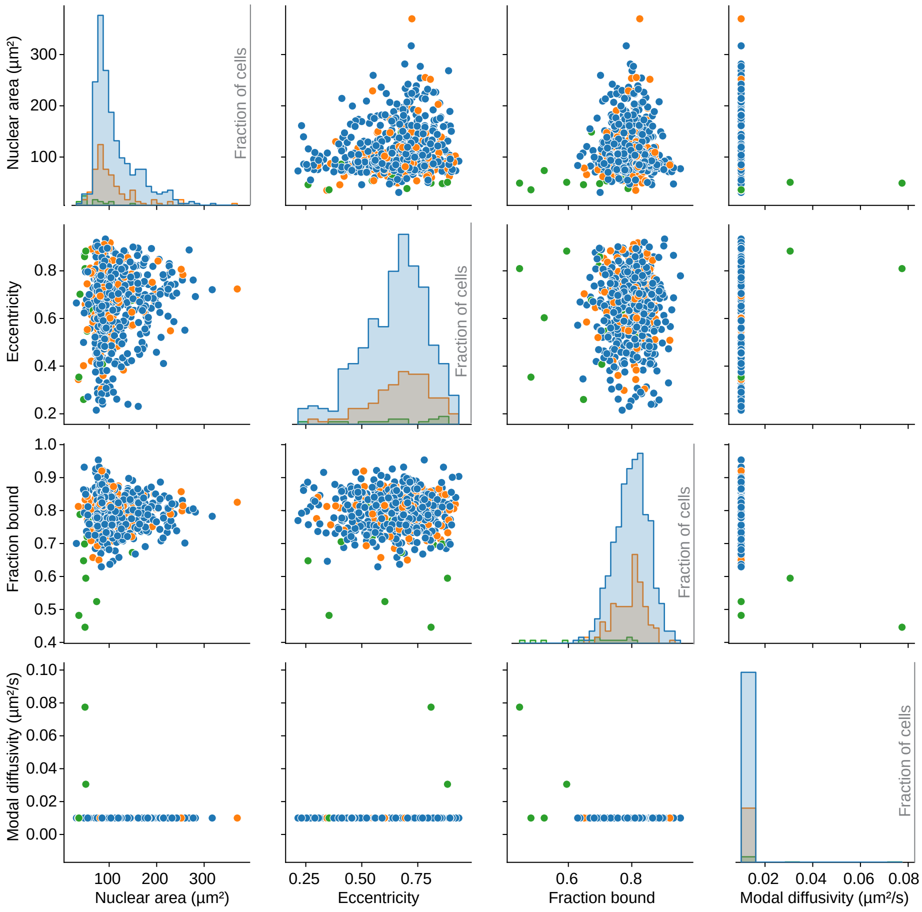

Figure S10

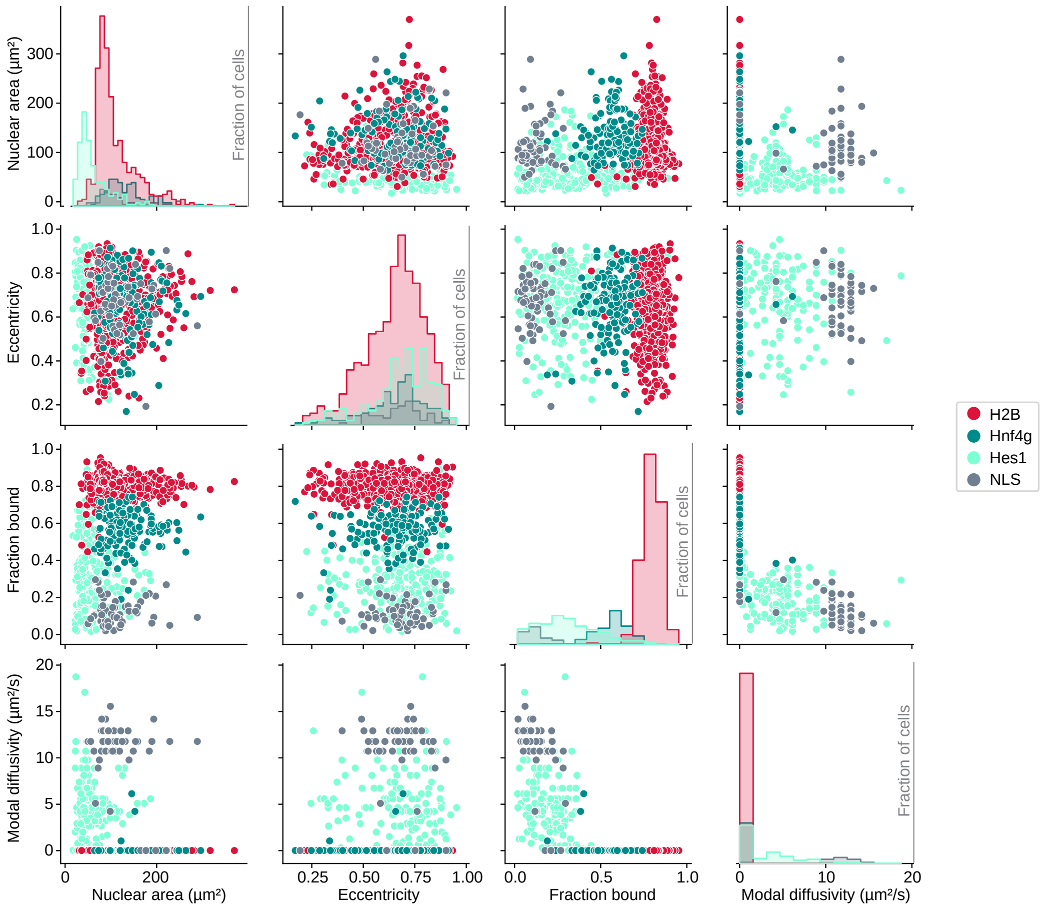
Nuclear area (µm²)
Eccentricity
Fraction bound
Modal diffusivity (µm²/s)
Fraction of cells
H2B
Hnf4g
Hes1
NLS

### Figure S11

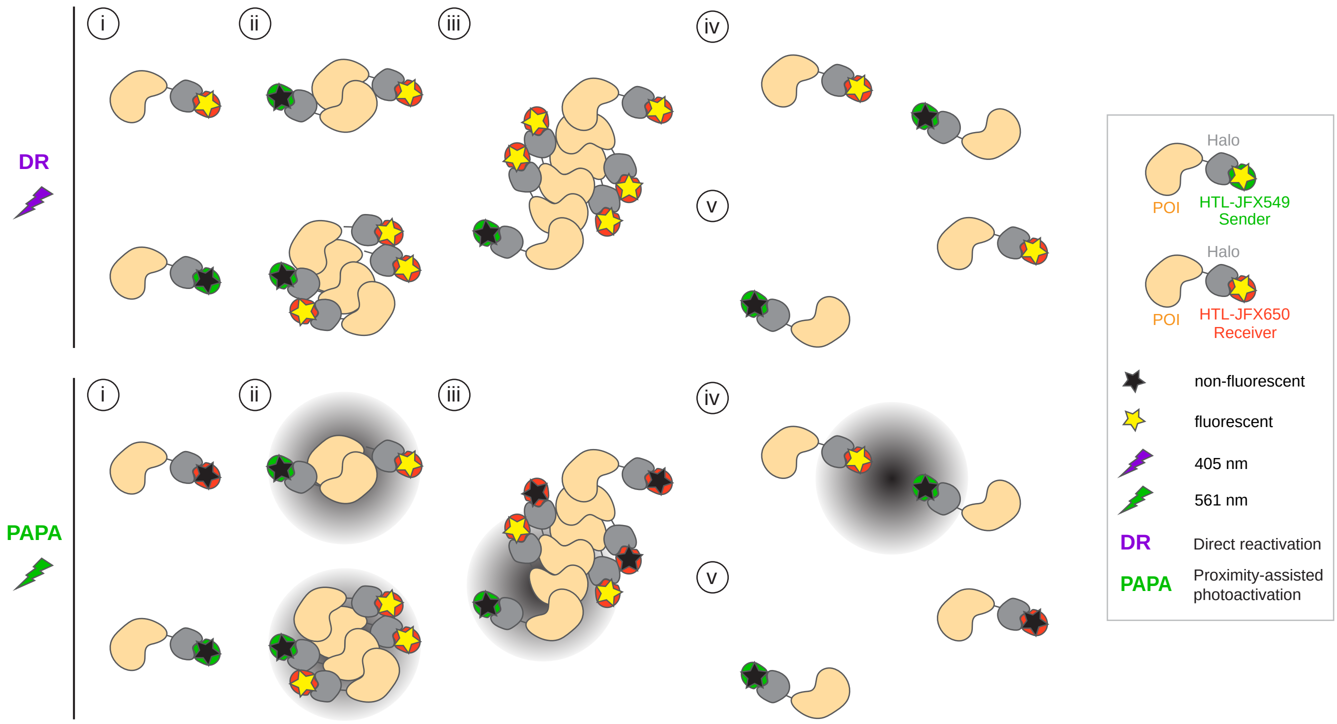
i
ii
iii
iv
v
DR
PAPA
Halo
POI
HTL-JFX549
Sender
HTL-JFX650
Receiver
non-fluorescent
fluorescent
405 nm
561 nm
DR
Direct reactivation
PAPA
Proximity-assisted photoactivation

### Figure S12

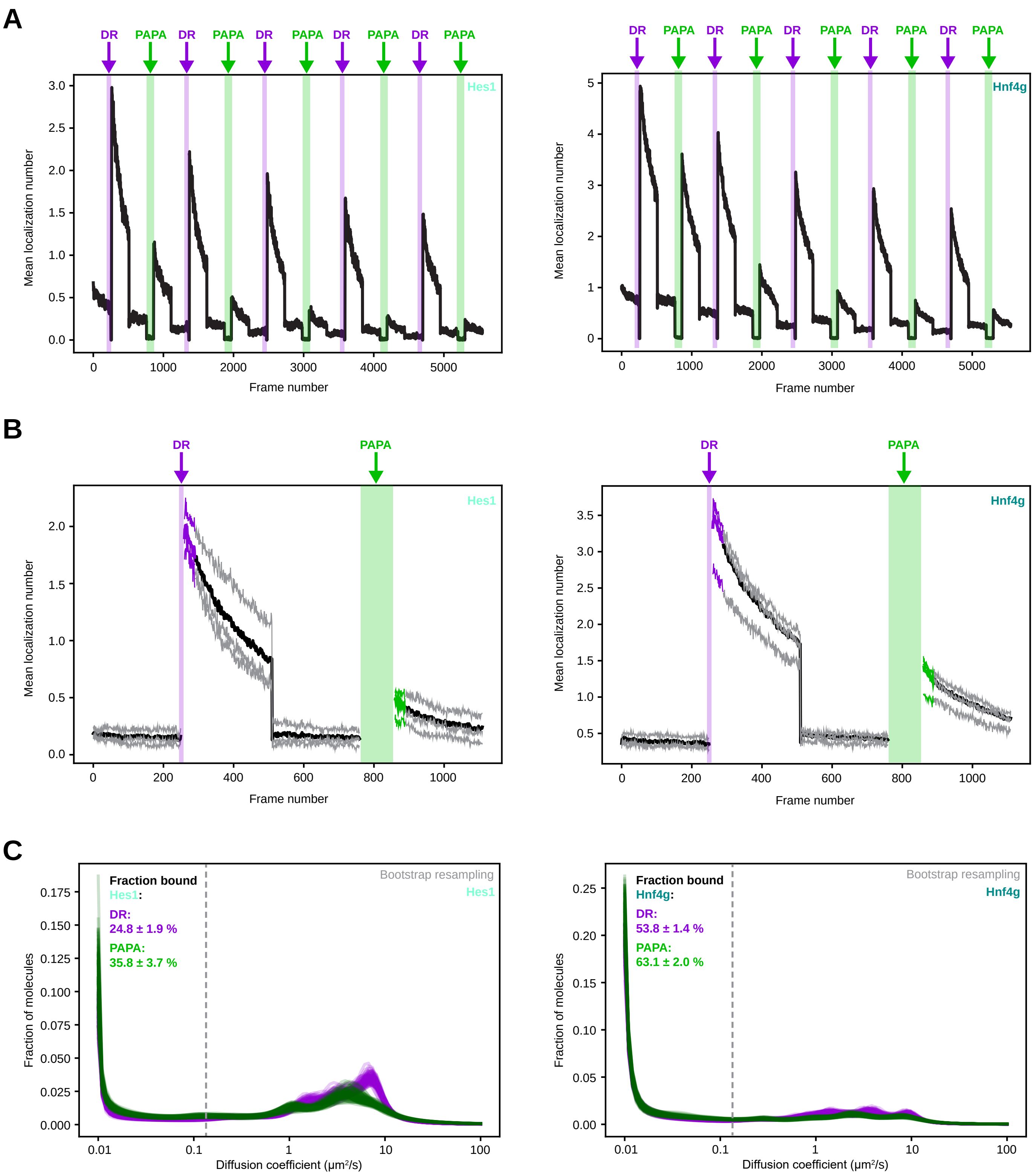
A
DR
PAPA
Hes1
Hnf4g
Mean localization number
Frame number
B
DR
PAPA
Hes1
Hnf4g
Mean localization number
Frame number
C
Fraction bound
Hes1:
DR:
24.8 ± 1.9 %
PAPA:
35.8 ± 3.7 %
Bootstrap resampling
Hes1
Fraction bound
Hnf4g:
DR:
53.8 ± 1.4 %
PAPA:
63.1 ± 2.0 %
Bootstrap resampling
Hnf4g
Fraction of molecules
Diffusion coefficient (μm²/s)
